# Quantification and Discovery of Acyl-ACPs by LC-MS/MS

**DOI:** 10.1101/870485

**Authors:** Jeong-Won Nam, Lauren M. Jenkins, Jia Li, Bradley S. Evans, Jan G. Jaworski, Doug K. Allen

**Author notes:** These authors contributed equally (Jeong-Won Nam, Lauren M. Jenkins). **One-sentence Summary:** Acyl-ACPs that enable fatty acid biosynthesis were uniquely profiled and quantified resulting in discovery of novel ACP products with unknown function and providing a method for rigorous comparisons. The author(s) responsible for distribution of materials integral to the findings presented in this article in accordance with the policy described in the Instructions for Authors (www.plantcell.org) is (are): Doug K. Allen and Bradley S. Evans.

## Abstract

Acyl carrier proteins (ACPs) are the scaffolds for fatty acid biosynthesis in living systems, rendering them essential to a comprehensive understanding of lipid metabolism; however, accurate quantitative methods to assess individual acyl-ACPs do not exist. A robust method was developed to quantify acyl-ACPs at picogram levels. Acyl-ACP elongation intermediates (3-hydroxyacyl-ACPs and 2, 3-*trans*-enoyl-ACPs), and unexpected medium chain (C10:1, C14:1) and polyunsaturated long chain acyl-ACPs (C16:3) were also identified, indicating the sensitivity of the method and that descriptions of lipid metabolism and ACP function are incomplete. Such ACPs are likely important to medium chain lipid production for fuels and highlight poorly understood lipid remodeling events in the chloroplast. The approach is broadly applicable to Type II FAS systems found in plants, bacteria, and mitochondria of animal and fungal systems because it uses a strategy that capitalizes on a highly conserved Asp-Ser-Leu-Asp (DSLD) amino acid sequence in ACPs to which acyl groups are attached. This allows for sensitive quantification using LC-MS/MS with *de novo* generated standards and an isotopic dilution strategy and will fill a gap in understanding, providing insights through quantitative exploration of fatty acid biosynthesis processes for optimal biofuels, renewable feed stocks, and medical studies in health and disease.

## INTRODUCTION

The synthesis of fatty acyl chains is essential for the production of storage and membrane lipids and functional molecules that modulate gene expression or contribute to protein activity. Studies on acyl chains span from disease and nutrition to demands for renewable fuels and feedstocks (Uauy et al., 2000; Peralta-Yahya et al., 2012; Zhu et al., 2016; Vanhercke et al., 2019). The lengths of up to 18 carbons are produced during fatty acid biosynthesis (Li-Beisson et al., 2013) on an acyl carrier protein scaffold (Overath and Stumpf, 1964) which is connected to the acyl chain through linkage of a 4’-phosphopantetheine group to a serine residue of the protein (ACP; Fig. 1a). The acyl chains are elongated by a series of four enzymatic steps that link ACP intermediates and form a cycle that results in the addition of two carbons and four hydrogens. Two distinct types of fatty acid synthase (FAS) complexes perform the repeated cycle of ketoacyl synthesis, reduction, dehydration, and a second reduction producing long chain saturated hydrocarbons (Fig. 1b). Yeast and mammals use a single large multifunctional FAS protein (Type I) that includes multiple domains for the enzymatic steps and ACP region (Bressler and Wakil, 1961; Hsu et al., 1965), whereas fatty acid synthesis in plants and bacteria occur through the concerted action of individual proteins within a complex including a distinct ACP that is 9-15 kDa in size (Alberts et al., 1963; White et al., 2005; Cronan, 2014).

**Figure 1.**
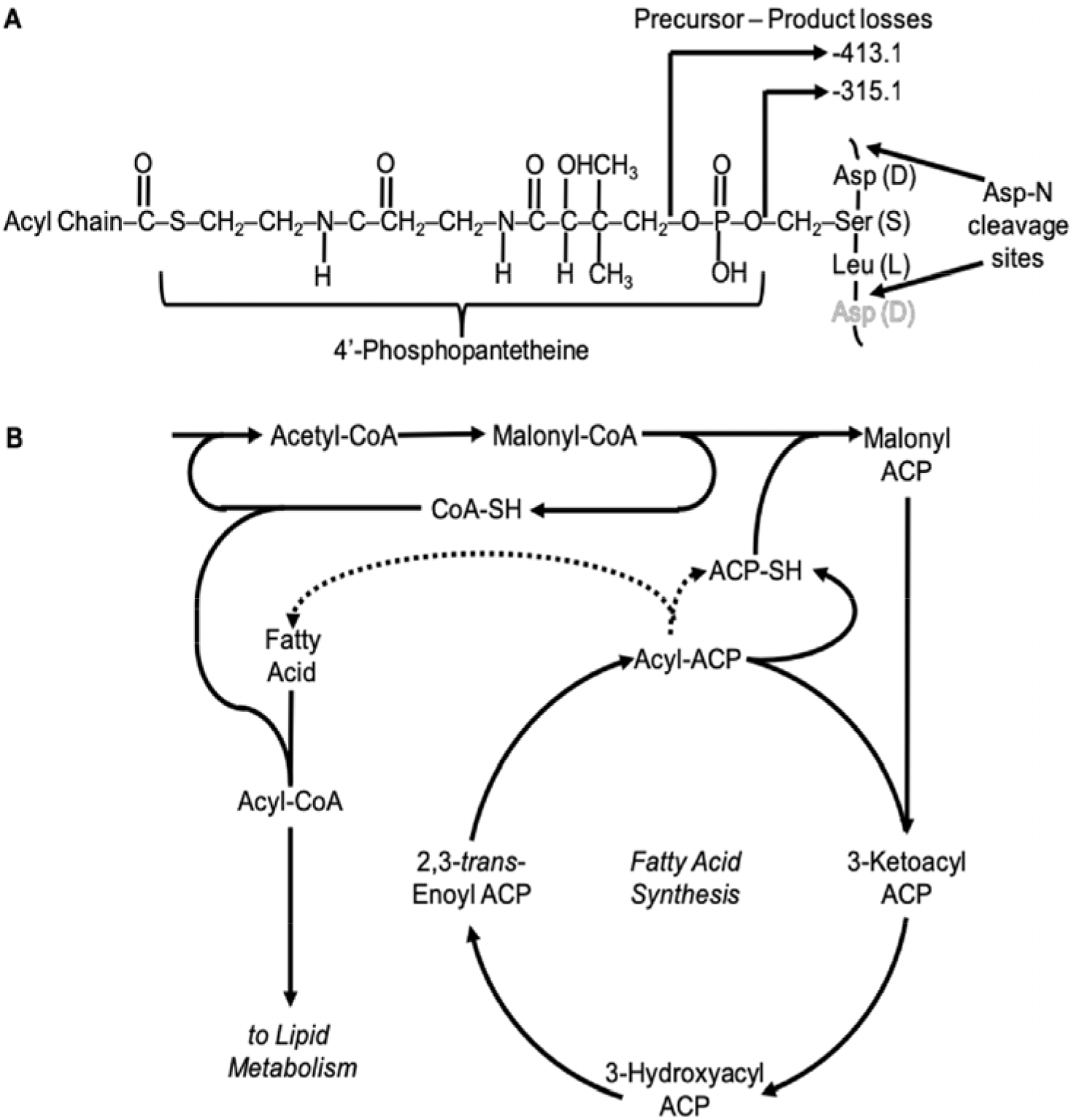
Acyl-ACP structure and function. **A)** Acyl-ACP is a 9-15 kDa protein with 4’phosphopantetheine arm attaching a fatty acyl chain to the protein backbone. The structure of acyl-ACP contains a highly conserved sequence of amino acids flanking the serine that attaches to phosphopantetheine. Peptide hydrolysis at aspartate residues produces an acyl molecular species that can be processed for characteristic fragments by mass spectrometry that have precursor-product losses of 315.1 and 413.1. **B)** ACPs are central to fatty acid synthesis with the process of fatty acid biosynthetic elongation of an acyl chain by two carbons taking place on ACP intermediates through a series of enzymatic steps that are repeated and conclude with hydrolysis by a thioesterase. The non-esterified fatty acid can then be used for lipid production.

In plants, the production of acyl chains take place predominantly in the chloroplast, though also to a lesser extent in the mitochondria, using acyl-ACP intermediates in both instances (Ohlrogge et al., 1979; Chuman and Brody, 1989). The chain elongation process is terminated by thioesterase reactions that release a non-esterified fatty acid from the ACP and regenerate the ACP substrate pool for further fatty acid biosynthetic reactions. The released fatty acid can then be activated to an acyl-CoA to make glycerolipids for storage oil production, waxes, surface lipids or membrane biogenesis (Li-Beisson et al., 2013).

Given the central role of acyl-ACPs in fatty acid biosynthesis, their quantitative analysis can help our understanding of lipid metabolism; however, the acyl group is a small percentage of the total acyl-ACP weight (usually significantly less than 3%) and current methods are confined by biochemical approaches that analyze intact proteins or their subunits. After ACP was demonstrated to be the acyl carrier for fatty acid synthesis (Goldman et al., 1963) studies examined expression (Hloušek-Radojčić et al., 1992; Bonaventure and Ohlrogge, 2002), subcellular roles (Brody et al., 1997; Wada et al., 1997; Cronan et al., 2005; Witkowski et al., 2007) and the impact of sequence on function and growth (De Lay and Cronan, 2007; Zhu et al., 2019); however studies quantifying the amount of acyl-ACPs are uncommon.

Levels of acyl-ACPs were measured based on urea polyacrylamide gel separation and immunodetection (Post-Beittenmiller et al., 1991); however multiple isoforms could not be resolved (Ohlrogge and Kuo, 1985) and only a subset of all acyl-ACPs were identified by antibodies and gel gradients, limiting the extent of quantification. Focusing more specifically on the acyl component, Kopka and coworkers (Kopka et al., 1995) developed an assay for individual acyl-ACP levels with gas chromatography linked to mass spectrometry (GC-MS) by generating butylamide derivatives. Levels of spinach leaf acyl-ACPs qualitatively matched those inferred from densitometry of urea-based gels; however, since the chemical derivatization was not specific to acyl-ACP thioester bonds, acyl-CoAs and other thioesters present at greater levels must be meticulously removed by anion exchange chromatography prior to derivatization. Moreover, short chain ACPs cannot be detected through this strategy.

We report a novel method to quantitatively analyze acyl-ACPs with tandem mass spectrometry based on a highly conserved contiguous amino acid region that flanks the attachment of the acyl chain. Enzymatic cleavage resulted in an acyl chain connected to a phosphopantetheinyl group and a tripeptide that was sensitively quantified by LC-MS/MS. ^15^N-labeled ACP standards were enzymatically synthesized *de novo* for absolute quantification of acyl-ACP levels using an isotopic dilution strategy. The method identified 3-hydroxyacyl-ACP and 2, 3-*trans*-enoyl-ACP, two of the three acyl-ACP elongation intermediates for each 2-carbon acyl-ACP product of the fatty acid biosynthetic cycle and unanticipated medium chain unsaturated acyl-ACPs that to date have no known function in plant metabolism. Tests with *Camelina sativa*, an emerging oilseed crop, indicated approximately 26 pmol/mgFW acyl-ACPs in developing seeds, present mostly in the terminal ACP product 18:1-ACP whereas leaves had much lower levels (approximately 3 pmol/mgFW). To our knowledge this is the only report of absolute quantification of resolved individual acyl-ACP levels in any organism and first application of a comprehensive approach to profile acyl-ACP products for improved understanding of fatty acid metabolism and lipid regulation.

## RESULTS

### Acyl-ACP standards were generated for absolute quantitation

Because acyl-ACP standards are not commercially available, isotopically labeled standards were generated using a 4’-phosphopantetheinyl transferase from *B. subtilis*, Sfp transferase (Sfp) (Fig. 2) which was overexpressed in *E. coli*. The apo-ACP substrate was also overexpressed in *E. coli* grown in LB media or ^15^NH_4_Cl to produce unlabeled and ^15^N-apo-ACP, respectively and the ACP products were purified by nickel affinity chromatography. Gel densitometry indicated the apo-ACP purity (79±5% SD, apo-ACP, n=36); with the remaining 21% phosphopantetheinylated. Protein concentration was calculated from absorbance and molar extinction coefficient. The unlabeled- and ^15^N-apo-ACPs (100-250 μM) and acyl-CoAs (300-500 μM) served as substrates that were enzymatically reacted with the Sfp transferase (25 μM) to produce individual acyl-ACPs.

**Figure 2.**
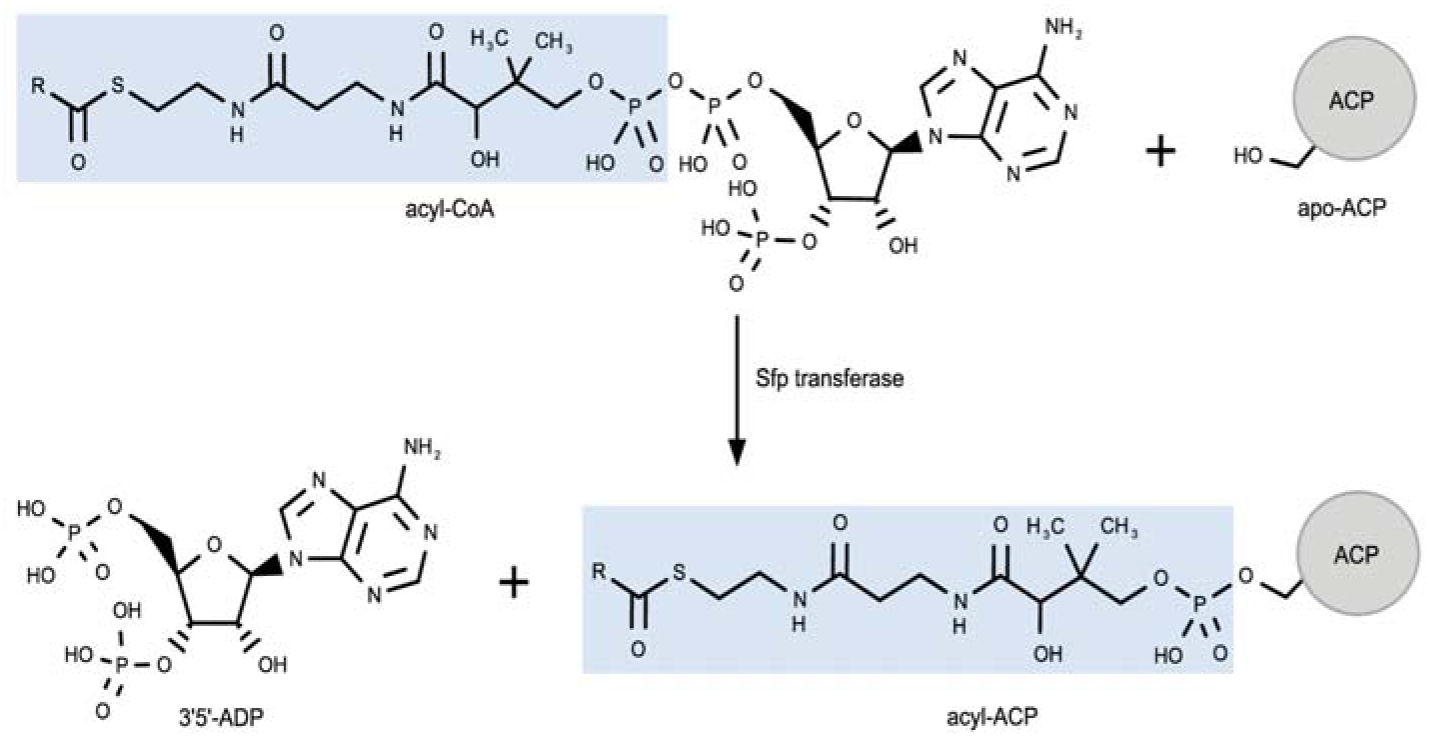
Enzymatic synthesis of acyl-ACP standards using Sfp transferase. Sfp transferase covalently transfers the acyl-4’-phosphopantetheine prosthetic group (highlighted in blue) from acyl-CoA to conserved serine sites on apo-ACP. Non-esterified (holo), saturated acyl chains with lengths of 2-18 carbons, and 16:1 and 18:1 acyl-ACP standards were synthesized. R, acyl-chains.

Acyl-CoA substrates with varying chain lengths were used to test the enzymatic reaction efficiency with varying buffer conditions and concentrations of dithiothreitol (DTT), MgCl_2_, and MnCl_2_. Buffering to pH 6.5 with 50 mM 3-(N-Morpholino)propanesulfonic acid (MOPS) improved product formation in agreement with prior research on phosphopantetheinyl transferases (Quadri et al., 1998; Sanchez et al., 2001; Joshi et al., 2003). Inclusion of DTT (1-5 mM) improved reaction efficiency in some instances but caused product degradation in others therefore 4 mM tris(2-carboxyethyl)phosphine (TCEP) was substituted to improve stability over a greater pH range and enhance disulfide selectivity. Additionally, 10 mM MgCl_2_ and MnCl_2_ were added to achieve near quantitative conversion of apo-ACP to short (C2) and medium chain (C8) acyl-ACP within a three-hour reaction time at 37°C and increase long chain (C18) reaction conversion, though not to completion (Supplemental Figure S1). The addition of MgCl_2_ and MnCl_2_ resulted in partial precipitation of medium and long chain acyl-CoAs (i.e. C10 – C18-CoA) (Constantinides and Steim, 1986) that was alleviated by DMSO (20% v:v) and TWEEN-20 (1% w/v) without impacting enzyme proficiency (Supplemental Figure S2). Using these conditions (Supplemental Table S1), yields greater than 90% were consistently achieved for all acyl chains apart from malonyl-ACP which could not be reliably obtained due to the unstable malonyl-CoA substrate that was easily decarboxylated.

### ACPs from many organisms share a conserved sequence that is efficiently cleaved with endoproteinase Asp-N

The elongating acyl chains are the component of interest within fatty acid synthesis; however, the acyl group is small in mass relative to the protein portion of acyl-ACPs. We hypothesized that methods to remove much of the protein chain would benefit quantitative analyses of the acyl composition. Although the amino acid sequence in ACP is diverse and varies between isoforms and across species, a highly conserved region fortuitously flanks the serine residue which joins the protein to the phosphopantetheine group. Vascular plant, algae, many bacteria, and mitochondrial ACPs from yeast, insects, and mammals, possess a four amino acid sequence (Asp-Ser-Leu-Asp; DSLD) surrounding the acyl attachment site (Fig. 1a, Table I). Therefore, proteolytic hydrolysis adjacent to aspartate residues with the protease Asp-N produces a three amino acid peptide linked to the 4’-phosphopantetheine and acyl group (Fig. 1a).

**Table I.**
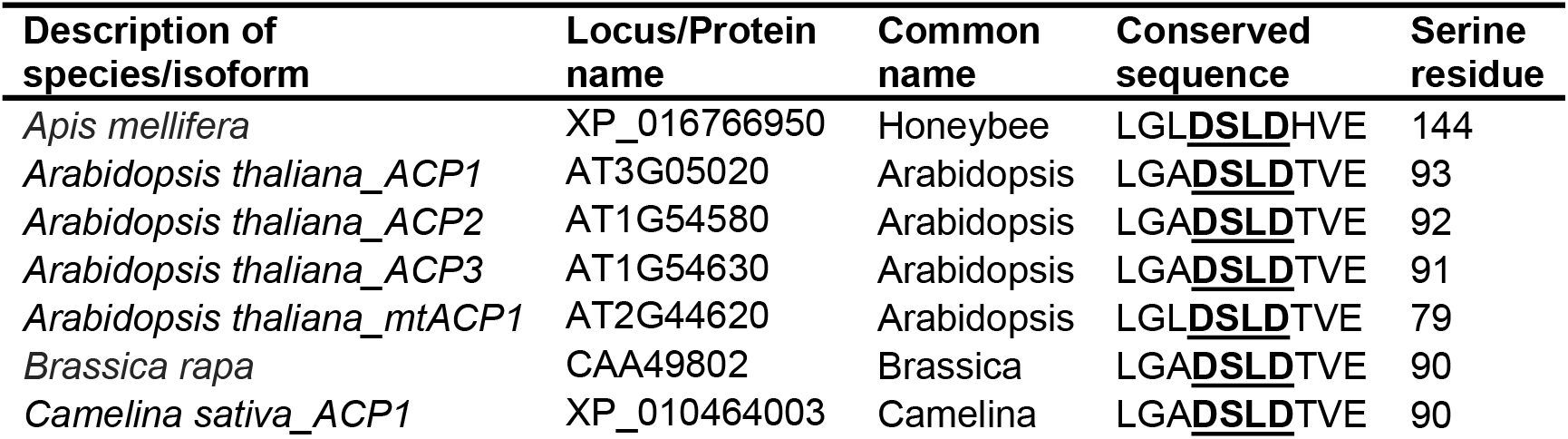
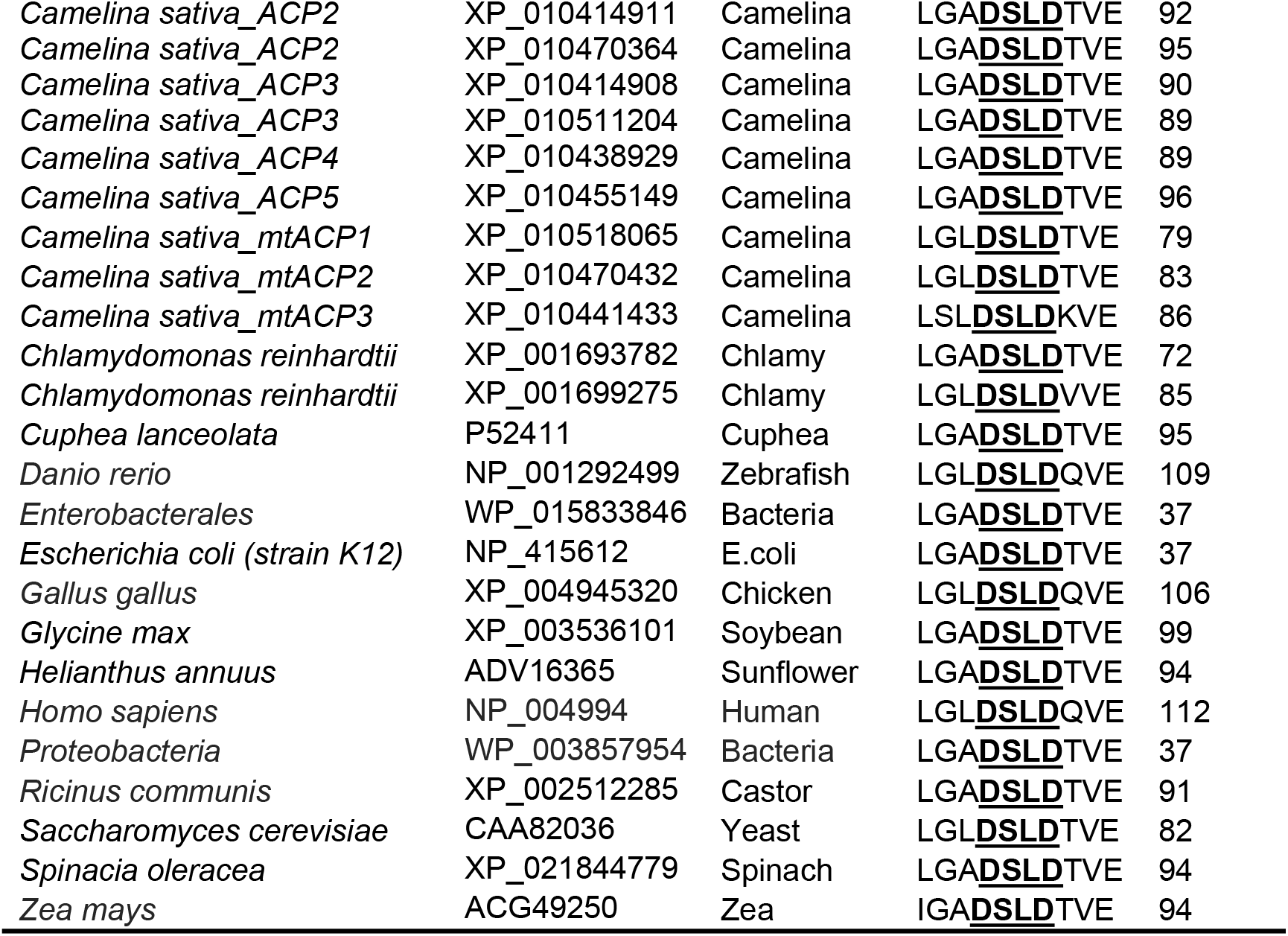
Highly conserved amino acid sequence in ACPs. Type II ACPs frequently contain a contiguous region of four amino acids (DSLD). The serine residue provides the point of attachment to the acyl limb described in Figure 1A. For most plant ACPs, only one chloroplast isoform is shown except where noted. Bacteria including: *E. coli* (strain K12), *Enterobacterales* and *Proteobacterial* sequences were inspected and yeast and animal mitochondria ACPs are also presented. Sequences for ACP protein alignment were obtained from NCBI protein database and are listed alphabetically by species.

ACP digestion was optimized using visual detection from the chromophore BODIPY-FL-*N-*(2-aminoethyl) maleimide-S-ACP (BODIPY-ACP) conjugate as described in the methods with several forms of Asp-N that are commercially available. Digestion of synthesized acyl-ACP standards was most efficient when the standards were purified away from other reaction components by trichloroacetic acid (TCA) precipitation, possibly due to substrate inhibition of Asp-N activity. Recovery of the acyl-ACP standards post-synthesis was assessed by gel densitometry (69±14% SD, n=9; Supplemental Figure S3). Acyl-ACP standards were digested with Asp-N endoproteinase at ratios of 1:20, 1:50, 1:100 (enzyme: acyl-ACP) at varying pH, temperature, and times. The reaction products were analyzed by SDS-PAGE to establish an adequate reaction system. Though short chain acyl-ACPs could be easily digested in less time and with less enzyme, obtaining complete digestion of medium-long chains required 1:20 ratio of enzyme to substrate for 16 hours at 37°C (Supplemental Figure S4). The digestions were quenched by the addition of methanol (50% of final concentration).

### Acyl-ACP digested products can be sensitively detected by LC-MS/MS

The digested acyl-ACP standards as well as biological samples were separated by reverse phase chromatography (C18) and analyzed by mass spectrometry in positive mode (i.e. *z*=1, *m/z* + 1 adduct), using a liquid chromatography tandem mass spectrometer as described in the methods (Fig. 3a). The masses calculated from chemical composition along with the multiple reaction monitoring (MRM) mode of analysis of the tandem MS were used to confirm the products of acyl-ACP fragmentation and optimize instrument performance. Two mass losses were consistently observed for all acyl-ACP digestion products indicating that they were not specific to the changing size in acyl chain, but instead components of the redundant 4’-phosphopantetheine and three amino acid sequence (Fig. 1a). A product ion that differed from the precursor by 315.1 indicated the loss of the three amino acids (DSL). The additional loss of phosphate increased the difference in mass to 413.1 and resulted in the primary product ion observed by mass spectrometry (Fig. 3b). Malonyl-ACP had an additional product ion with a difference in mass of 457.1, that indicated the loss of the DSL, phosphate, and the carboxyl from the malonyl-group and is the primary product ion for this molecule. The ^15^N labeled standards are 3 Da heavier giving a precursor-product ion difference of 318.1 and 460.1. The changes in abundance were used to optimize the declustering potential (DP), collision energy (CE) and collision cell exit potential from standards and acyl-ACPs isolated from plant biomass (Fig. 3c). Optimal chromatography and detection on the mass spectrometer of acyl-ACP digested products was indicated by spectra with well-defined peaks that were also evenly spaced to represent the change in acyl mass by consistent increments of C_2_H_4_. All acyl species from acetyl-ACP to 18 carbon acyl chains (i.e. 18:0-ACP) were identified in addition to non-esterified (i.e. holo) and apo-ACP. The dynamic range (approximately 3-4 orders of magnitude) was established through comparison of integrated peak areas for each acyl-ACP, with the exception of holo- and malonyl-ACP due to dimerization and instability, from standards and Camelina samples and indicated that femtomole (i.e. picogram of protein) quantities could be quantified accurately (Fig. 3c, Supplemental Figure S5). Since pooled acyl-ACPs are found at low concentrations (10 ng/mg FW in spinach chloroplast (Ohlrogge et al., 1979)), saturation of the detector is unlikely to be an issue for most preparations.

**Figure 3.**
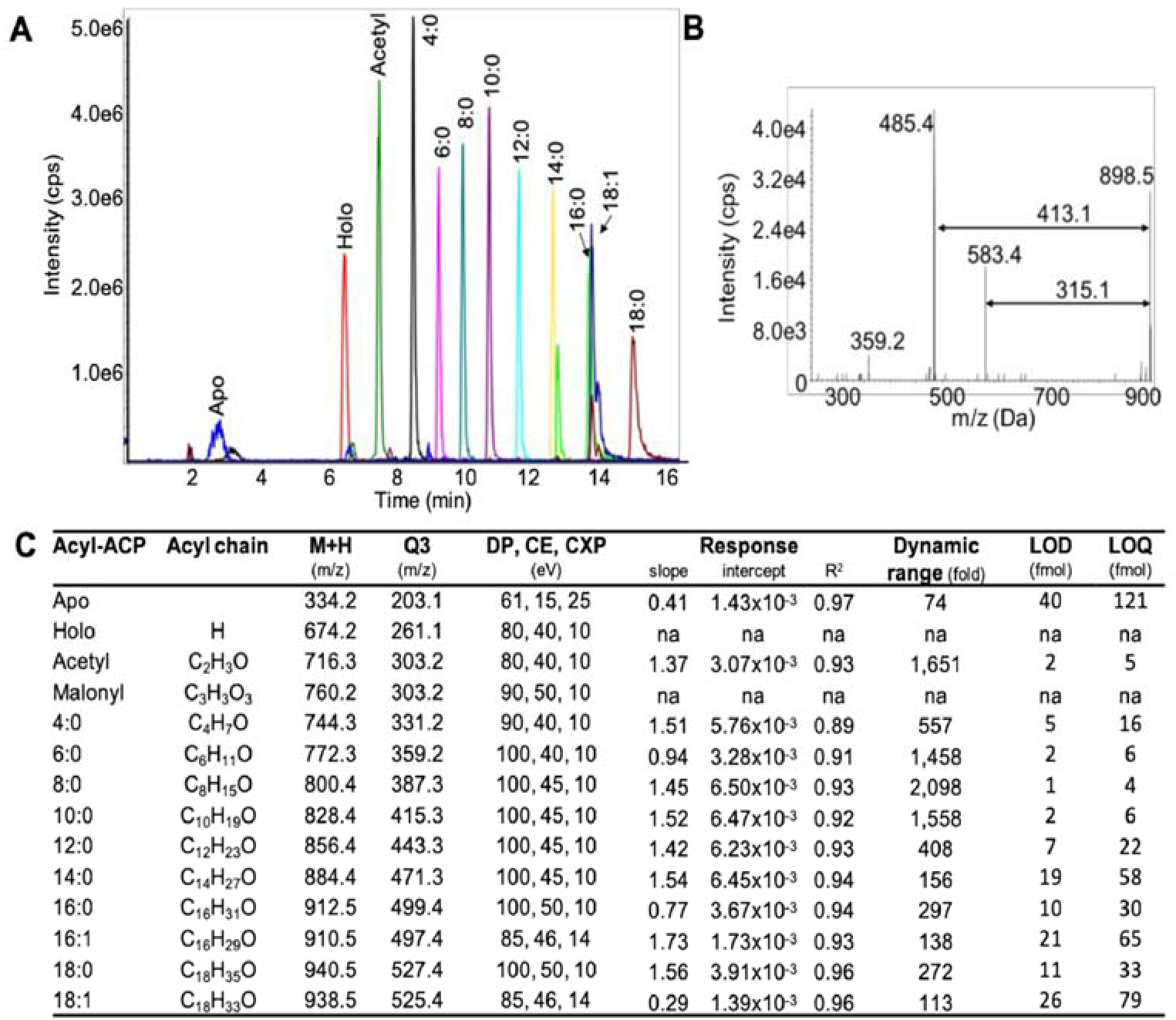
Mass spectral detection and analysis of acyl-ACPs. **A)** Chromatographic properties of acyl-ACP standards. Asp-N digested partial acyl-ACPs are separated according to the length and properties of the acyl chain on a reversed phase column by liquid chromatography and detected by triple quadrupole mass spectrometer. Labels above peaks represent the acyl species in the acyl-ACP molecules. **B)** MS/MS fragmentation product ion scan of Asp-N digested 15:0-ACP standard indicating the major losses. **C)** Optimal mass spectrometry parameters for acyl-ACP detection by multiple reaction monitoring (MRM). Declustering potential (DP), collision energy (CE), collision cell exit potential (CXP) Responses are represented as the slope (ratio of peak areas per pmol analyte) of the calibration curve for each ACP species. Linear regression (R^2^) weighted by 1/Y for all samples. The standard deviation of the lowest detectable levels and slope of the calibration curve were used to establish the limit of detection (LOD) and quantification (LOQ).

### Total acyl-ACP levels in Camelina seeds are approximately 17-fold greater than leaves

Isotope dilution, a general strategy which relies upon internal standard addition for derivation of calibration curves, is the benchmark for quantitation by mass spectrometry. Isotopically labeled internal standards provide chemically equivalent ions distinguished from sample metabolites, or analytes, by a change in mass corresponding to the isotope label(s) (in this case the addition of three ^15^N atoms in place of ^14^N shift the mass by ~3 Da). The addition of internal standards at an early stage of sample preparation allows one to normalize data and rectify sample to sample variations in analyte recovery. In addition, the chemical equivalency of isotopically labeled standards can account for differences in observed analyte response factors and address the exposure to differential matrix effects. Thus, an isotope dilution strategy is superior to other approaches that rely on standards that are not chemically equivalent with differences in response factors and subject to different matrix effects amongst analytes that are not accounted for resulting in inaccuracies. Here, an isotope dilution strategy was used to quantify acyl-ACP levels in Camelina seeds and leaves. Analyte acyl-ACPs from plant biomass were spiked with internal standards before extraction and then quantified. Total ACP amounts detected in seeds were ~26 pmol/mgFW composed predominantly of 18:1-ACP (Fig. 4). Camelina leaves exhibited ~9-fold lower levels overall than seeds; consistent with calculated estimates from the literature (~1-2 pmol/mgFW calculated from spinach leaves (Ohlrogge et al., 1979)), and of comparable composition (Post-Beittenmiller et al., 1991).

**Figure 4.**
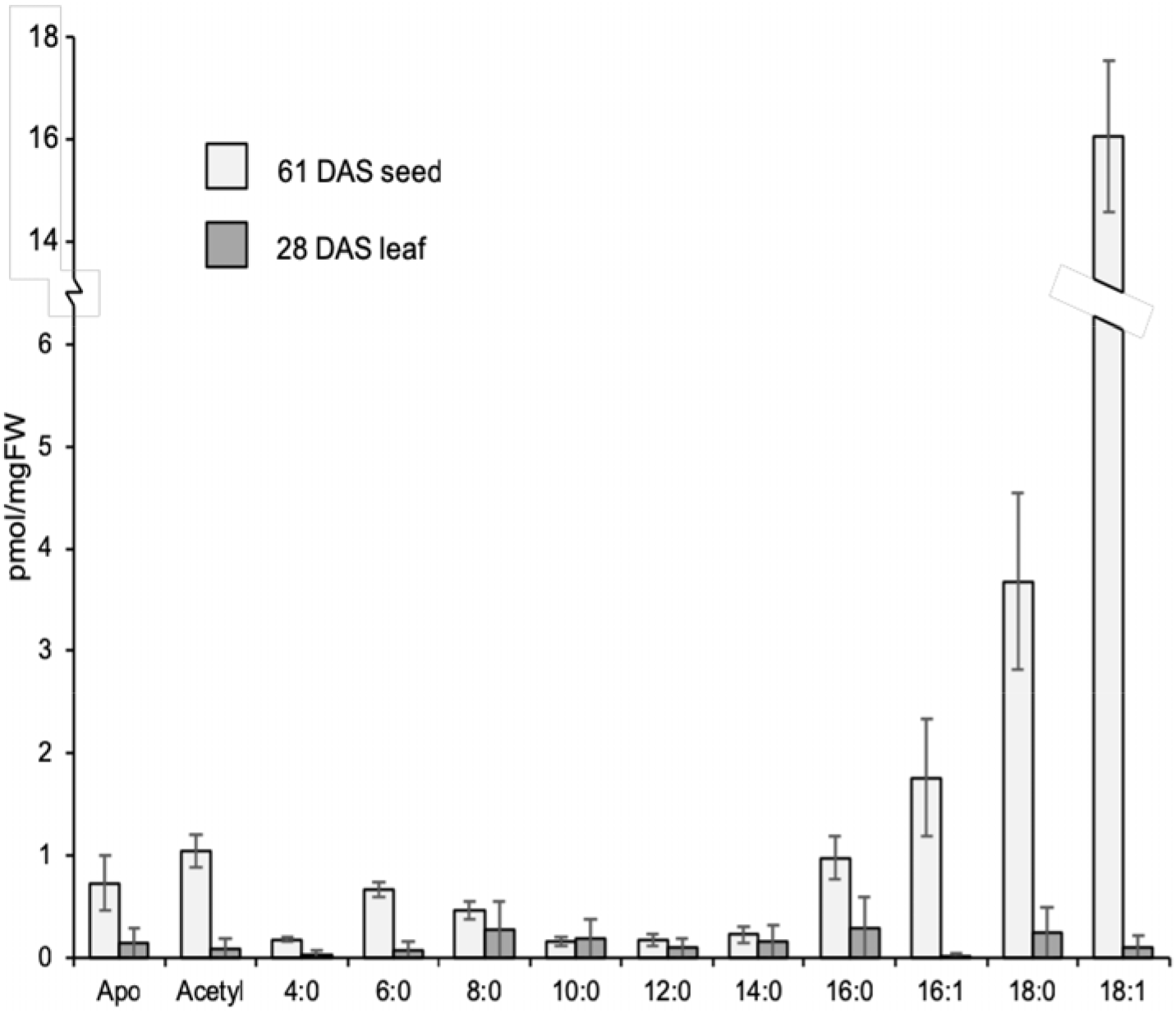
Acyl-ACP levels in Camelina seed and leaf. Acyl-ACPs from 61 DAS seed (n=11-15; except 16:1, 18:0, and 18:1 n=4) and 28 DAS leaf (n=16-19; except 16:1, n=4; 18:0, n=6; 18:1 n=12) were quantified using internal standards. Lower replication of saturated and unsaturated long chains reflected initial digestion inefficiencies in sample preparation. The means and standard deviations are presented for each acyl species.

### Two acyl-ACP intermediates of the fatty acid biosynthetic cycle were identified and previously undescribed unsaturated acyl-ACPs were discovered

The process of acyl chain extension in fatty acid synthesis is completed by a cycle that generates three additional acyl-ACP elongation intermediates reflecting the ketoacyl condensing, reducing, and dehydrating steps before the final reduced acyl-ACP. Though these intermediates are believed to be present at low quantities we considered the possibility of their detection. Since the intermediates are themselves ACP products that differ only in the acyl structure and composition, the developed methods were presumed applicable though potentially limited by the instrument sensitivity.

Analysis of seeds indicated several additional acyl-ACP peaks. We calculated the molecular weights for the elongation intermediates assuming a loss of 413.1; identical to other acyl-ACPs measured. Two acyl-ACP elongation intermediates were detected for each acyl chain-length form: 3-hydroxyacyl-ACPs and 2, 3-*trans*-enoyl-ACPs. The MRM list (Supplemental Table S2) also included the exact masses for 3-ketoacyl-ACPs, the first intermediate in the fatty acid biosynthetic cycle (Fig. 1b); though this intermediate was not detected. The 3-hydroxyacyl-ACPs in developing seeds were present at similar amounts to most reduced acyl-ACPs (Fig. 5a) with heightened levels of C14-hydroxyacyl-ACP that may be involved in Lipid A-like molecule production in the mitochondria (Wada et al., 1997; Li et al., 2011). Peaks for the 2, 3-*trans*-enoyl-ACP were lower in abundance, generally less than 2.0% of total (C18-enoyl; Fig. 5a).

**Figure 5.**
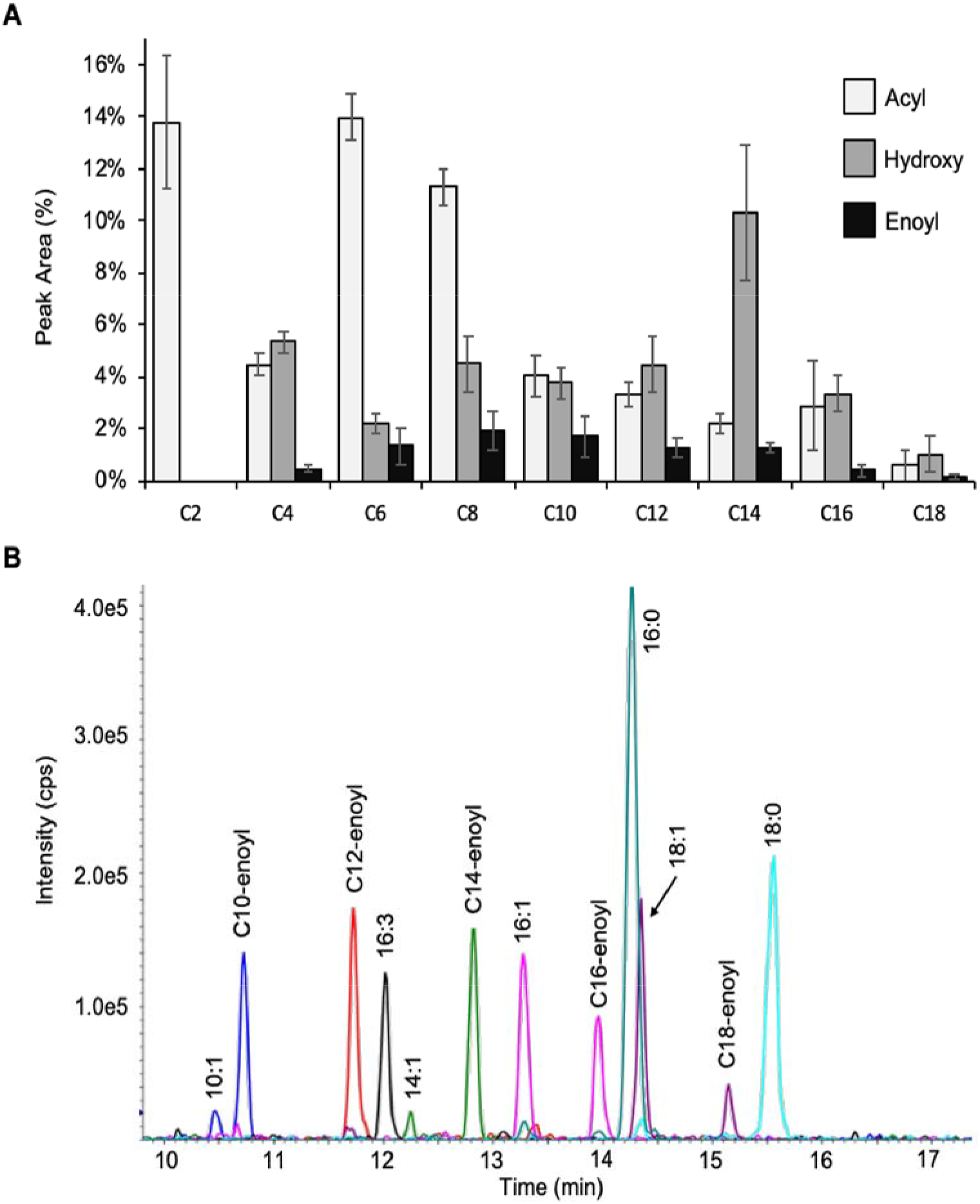
Acyl-ACP elongation intermediates detected from Camelina developing seeds. Results (means and standard deviations) from a liquid chromatography tandem triple quadrupole mass spectrometer analysis. **A)** Relative peak areas in percentage were calculated from all the acyl-ACP species detected. The average and standard deviation are presented based on four independent sample preparations. **B)** Chromatographic detection of unanticipated unsaturated acyl-ACPs. The characteristic double peak pattern for the 2,3-*trans*-enoyl- and desaturated acyl-ACP isomers were observed for two medium chain acyl-ACPs (C10, dark blue; C14, green). Polyunsaturated C16-ACP (16:3-ACP; black) was also detected.

The 2, 3-*trans*-enoyl-ACP and equivalent chain length single desaturated acyl-ACPs are isomeric, therefore, for example, 16:1-ACP (*cis*-double bond) and C16-enoyl-ACP (2, 3-*trans-*double bond) cannot be discriminated by mass. Consequently, acyl chain MRM transitions indicated a second peak, the 2, 3-*trans*-enoyl acyl-ACP, at a different retention time. To conclusively identify the isomers, a standard of 18:1-ACP was digested and processed resulting in a peak that eluted at the same retention time as one of the two peaks in question (Supplemental Figure S6). This confirmed the designation of each peak, with the 2, 3-*trans*-enoyl acyl-ACP characteristically eluting after the *cis*-unsaturated acyl-ACP as a much smaller peak. The 3-hydroxyacyl-ACP products that differ in mass eluted ahead of the corresponding acyl-ACP as presented in a summary of the retention times in Supplemental Table S3.

Interestingly, owing to the sensitivity of the method, we observed peaks corresponding to several medium chain unsaturated acyl-ACPs (i.e. C10:1 and C14:1) and a significant amount of polyunsaturated acyl-ACP (16:3) in seeds and leaves (Fig. 5b) that have not been previously described. The paired MS/MS transitions, retention times, and peak shapes were all consistent with expectation for these species supporting their identification. The retention time offset (Fig. 5b) was similar to measured differences for 9-*cis*-18:1-ACP (Fig. S6) and 7-*cis*-16:1-ACP and indicated that these unsaturated forms likely contain *cis* double bonds. Enoyl-ACPs contain *trans* double bonds and have a near-linear predicted 3D structure and therefore more closely approximate saturated ACPs with respect to their physical properties. *cis*-unsaturated-acyl groups are commonly generated within living systems, though are not believed to be biosynthetic intermediates on fatty acid synthase assembly lines; thus, the roles in metabolism for these molecules remains to be elucidated.

## DISCUSSION

### Development of an acyl-ACP profiling method based on a conserved contiguous peptidyl region

Acyl chains represent 90-95% of the carbon in glycerolipids and thus the production and movement of fatty acids is central to lipid metabolism. We developed a sensitive method to identify and quantify acyl-ACPs that can complement analysis of acyl-CoAs providing a more complete picture of the connectivity between fatty acid biosynthesis and lipid assembly (Allen, 2016a). The growing and nascent acyl chains are tethered to the ACP via a 4’-phosphopantetheinyl linker and to a serine residue that is part of a conserved four amino acid phosphopantetheine attachment site (Asp-Ser-Leu-Asp; DSLD). Acyl-ACPs are present in cells in scant quantities and are made from different ACP isoforms. Each can be reduced with an aspartyl protease to three amino acid peptides linked to 4’-phosphopantetheine that differ only in the attached acyl chains. This effectively eliminates protein size and sequence variation that convolute the population of a single acyl-chain species across multiple possible carriers and enables their detection in forms that are convenient for mass spectrometric analysis.

In comparison to described prior attempts the digestion and LC-MS/MS-based analysis does not require antibodies, multiple urea-PAGE gels, or derivatization techniques and requires only mg levels of biological material. Acyl-ACPs that are present at picogram/mgFW quantities can be readily detected without interference from acyl chains esterified to lipids or CoA species that are more abundant in biomass. In addition, the use of the phosphopantetheinyl transferase Sfp, from *B. subtilis*, which tolerates a wide variety substituents added onto its canonical substrate, 4’-phosphopantetheine, enabled a one-step synthesis of acyl-ACPs starting with acyl-CoAs and ACP (Fig. 2) (Lambalot et al., 1996). The approach circumvented a more typical, two step enzymatic synthesis beginning with production of holo-ACP, often with the E. coli phosphopantethienyl transferase AcpS, followed by acylation with acyl-ACP synthetase (AAS) to generate acyl-ACPs (Zornetzer et al., 2006).

### Acyl-ACP profiling reveals unanticipated acyl-ACP groups with implications for lipid metabolism in oilseeds

We quantified the absolute levels of individual acyl-ACPs containing up to 18 carbons for the first time. 18:1-ACP was present in the greatest quantities in seeds and may indicate fatty acid biosynthesis is not a bottleneck for lipid production in oilseeds but could be important to regulation (Andre et al., 2012). In addition, two intermediates of the fatty acid biosynthetic cycle, 3-hydroxyacyl-ACP and 2,3-*trans*-enoyl-ACP were identified. Interestingly, our approach elucidated other unsaturated acyl-ACPs in addition to 18:1, including medium chain (C10:1 and C14:1) that have unknown function and that have not been previously reported. Speculatively, it would be interesting to determine whether plants or other species that have enhanced medium acyl chain lipid production also have higher levels of these ACP-forms that could indicate differences in the fatty acid biosynthetic machinery which would be targets for metabolic engineering. Higher degrees of unsaturation in C16 acyl chains (i.e. 16:3-ACP) were also observed (Fig. 5) which may be relevant to lipid remodeling in the chloroplast. The detection of unanticipated acyl-ACPs indicates that the method can serve as a sensitive and quantitative tool for discovery aspects of fatty acid metabolism.

### Absolute levels of Acyl-ACPs are greater in seeds than leaves of oilseeds

The levels of acyl-ACPs in seeds were greater than leaves. Oilseeds actively produce storage lipids that can represent the largest fraction of seed biomass whereas leaves make a small amount of lipids primarily for use in membranes (Fan et al., 2014). Similar to more qualitative studies on spinach leaves (Post-Beittenmiller et al., 1991), our method quantified high levels of long chain acyl-ACPs (Fig. 4a), predominantly 18:1, in seeds. The present method also detected an apo-ACP pool in camelina seed/leaf that was not monitored in the spinach studies. The 18:1-ACP is a terminal product of fatty acid synthesis and may indicate that acyl chain production is not limiting lipid biosynthesis. The presence of an apo-ACP pool could also suggest that ACP abundance is not limiting fatty acid production, but perhaps the activation to holo-form and the source of acetyl groups impact the regulated rate of the fatty acid biosynthetic cycle. Future studies can potentially capitalize on this method through transient isotopic labeling studies (Allen, 2016b) to assess synthetic and turnover characteristics in lipid metabolism more rigorously.

### The role of acetyl-ACP in plant lipid metabolism and regulation

In spinach leaves and seeds, 5-10% of the total ACP is in an acetylated form (Post-Beittenmiller et al., 1991) which is comparable to numbers reported here for Camelina tissues (4% and 11% for seed and leaf, respectively; Fig. 4). *In vitro* studies have previously indicated that acetyl-ACP is inefficiently but preferentially used by KAS I (as opposed to KAS III) condensation reactions (Jaworski et al., 1993) and it can also be converted to acetyl-CoA by the reversible acetyl-CoA: ACP transacylase reaction. Given the reversibility of transacylase reactions, the steady state levels of acetyl-ACP may reflect demands for acetyl-CoA (Post-Beittenmiller et al., 1991) and regulate acetyl-CoA carboxylase (Post-Beittenmiller et al., 1991; Andre et al., 2012). In the future, studies on lipid metabolism will benefit from a higher resolution understanding of the fatty acid biosynthetic cycle and regulation of its elongation intermediates which can now be accurately quantified.

## MATERIALS AND METHODS

### Chemicals, media, reagents and materials

All chemicals and media were from Sigma (St. Louis, MO) unless otherwise noted. Coenzyme A, malonyl-CoA, acetyl-CoA, butyryl-CoA, hexanoyl-CoA, octanoyl-CoA, decanoyl-CoA, lauroyl-CoA, myristoyl-CoA, were from Sigma. Palmitoyl-CoA, stearoyl-CoA, 16:1(n7)-CoA, 18:1(n9)-CoA were from Avanti Polar Lipids (Alabaster, Alabama). BODIPY-FL-N-2-aminoethyl)maleimide (BODIPY), Tris(2-carboxyethyl)phosphine (TCEP) and Gelcode stain was from Thermo Fisher (Waltham, MA). 14:0, 15:0, and 16:0-ACPs when the project was initiated were gifts from Dr. Ed Cahoon (University of Nebraska-Lincoln). SDS-PAGE and agarose gel supplies and Protein Assay Reagent were from Bio-Rad (Hercules, CA). Dimethylsulfoxide (DMSO) was from Chem-Impex International (Wood Dale, IL). The 5 mL HisTrap™ column and the 5 mL HiTrap™ desalting column used for protein purification were from GE Healthcare (Pittsburgh, PA). Amicon^®^ Ultra centrifugal filters used for protein concentration and thin layer chromatography (TLC) Silica gel 60 plates were from Millipore Sigma (St. Louis, MO). Restriction enzymes, T4 DNA ligase and DNA polymerase (Finnzymes’ Phusion-Hot Start) were from New England Biolabs (Ipswich, MA). Growth media were from BD Biosciences (San Jose, CA). Oligonucleotides were purchased from Integrated DNA Technologies (Coralville, IA) and listed in Supplemental Table S4. PCR cleanup, plasmid miniprep and gel extraction kits were from Qiagen (Valencia, CA). Antibiotics were used at the following concentrations: ampicillin or carbenicillin 100 μg/mL, kanamycin 50 μg/mL. *Camelina sativa* plants were grown in the Donald Danforth Plant Center greenhouse. Greenhouse conditions: temperature, day/night 21°C/20°C; humidity, 40%; light/dark 14h/10h.

### Microorganisms

*E. coli* DH5α λpir was a gift for William Metcalf and was used for cloning and plasmid maintenance. *E. coli* BL21 was purchased from New England Biolabs and was used for protein overexpression. E. coli BL21 (DE3) was purchased from Novagen (Danvers, MA) and used for protein over expression. *E. coli* BW25113 was obtained from as part (parent strain) of the Keio collection (Coli Genetic Stock Center, Yale University). Microorganisms were maintained either on agar selection plates for short-term storage or as a concentrated cell suspension in a 50% medium-glycerol slurry flash frozen in liquid nitrogen and kept at −80° C for long-term storage. DNA transformations were by electroporation of homemade electrocompetent cells.

### Plasmids

Plasmid pQE-60-eGFP was constructed by ligation of empty pQE-60 vector (Qiagen) and plasmid pEGFP-NI (clontech) both of which had been double digested with NheI and XhoI followed by gel purification.

Plasmid pT5Kan (Supplemental Figure S7a) construction began with digestion of pQE60 (Qiagen) with XhoI and NheI which had previously been modified to contain *eGFP* (pQE60-eGFP (unpublished)) at the NcoI/BamHI sites of the MCS. The gel-purified fragment was ligated to the PCR product of pET-28a (Novagen) amplified by primers pET-28-XhoI-F and pET-28-NheI-R which had been similarly digested and gel purified. The cloned product resulted in an intermediary plasmid that was further modified to remove unnecessary intergenic regions in order to make the vector easier to work with. The truncation was accomplished by PCR amplification of the intermediary plasmid using primers pT5-Kan-NotI-F and pT5-Kan-NotI-R which amplified the lacI-MCS-KanR-ColEI region leaving only 300 bp separating the *lacI* and ori-ColEI sequences as well as provided a NotI site for digestion and ligation of the amplified fragment. The cloned product, named pT5-Kan was used for cloning and expression of the *E. coli* acyl carrier protein gene, *acpP*.

Plasmid pT5-Kan-AcpP-His_6_ was constructed from the PCR product amplified from genomic DNA isolated from *E. coli* MG1655 using primers Acp-His_6_-F and Acp-His_6_-R and after digestion with NcoI/BamHI followed by gel purification was ligated to similarly digested and purified pT5-Kan.

Plasmid pET-Duet-1-Sfp was constructed from NdeI/KpnI digested PCR product amplified from pUC-18 Sfp (a gift from Christopher T. Walsh(Lambalot et al., 1996)) using primers Sfp-F and Sfp-R.

Plasmid pET-28a-Sfp-His_6_ (Supplemental Figure S7b) was constructed from NcoI/BamHI digested and gel purified PCR product amplified from pET-Duet-1-Sfp using primers Sfp-His_6_-F and Sfp-His_6_-R and similarly digested and purified pET-28a.

### DNA Sequencing and analysis

Plasmids were sequenced by Sanger sequencing on an ABI 3730XL capillary sequencer (University of Illinois Biotechnology Center) or ABI 3500 Genetic Analyzer (St. Louis Community College BioBench CRO Facilty) on linear amplification reaction products generated using BigDye Version 3.0 terminator/enzyme mix and standard protocols. Sequence outputs were assembled and analyzed using Sequencher 4.6 software (Gene Codes, Ann Arbor, MI). The primers used in the BigDye reactions were the same as were used for cloning except for the primers used for sequencing the acpP-His6 construct which used primers pT5-Kan-F and pT5-Kan-R.

### Overexpression and purification of apo-ACP and Sfp transferase

*AcpP from E. coli* and Sfp from *B. Subtilis* were expressed as C-terminal hexahistidine fusion in appropriate *E. coli* expression strains. For unlabeled protein preparations, the expression strains were grown in lysogeny broth (LB) supplemented with kanamycin (50 μg/mL) starting from 1% inoculations from overnight pre-cultures. The cells were grown at 37 °C on an orbital shaker (200 rpm) until mid-log phase (OD_600_ = ~0.5-0.8) before shifting the temperature to 18° C and addition of IPTG to a final concentration of 1 mM for overnight induction of expression. Isotopically labeled AcpP was expressed in a modified MOPS minimal medium containing ^15^N-labeled ammonium chloride (9.69 mM or 528 mg/L) and kanamycin (50 μg/mL); growth temperature and induction conditions were the same as for the unlabeled preparations. Cells were harvested by centrifugation, resuspended in lysis buffer (10 mM imidazole, 20 mM HEPES, 150 mM NaCl, 1 mM dithiothreitol (DTT), 10% glycerol, 10 mg/mL lysozyme, protease inhibitor), and lysed by sonication: 6 cycles of 1 minute 50% duty cycle followed by 1 minute cooling time at 4° C using a micro-tip probe sonicator. The lysates were then clarified by centrifugation for 20 minutes at 20,000 rpm before purification. The clarified extracts were applied to a 5 mL HisTrap column (GE Healthcare) using an AKTA™ FPLC (GE Healthcare), washed with 20 column volumes of loading/wash buffer (10 mM imidazole, 20 mM HEPES, 150 mM NaCl, 1 mM DTT, 10% glycerol) before elution with elution buffer (250 mM imidazole, 20 mM HEPES, 150 mM NaCl, 1 mM DTT, 10% glycerol). Target protein containing fractions were concentrated by ultrafiltration using Amicon® Ultra Centrifugal Filters (Millipore Sigma) before desalting and buffer exchange into storage buffer (20 mM HEPES, 150 mM NaCl, 1 mM DTT, 10% glycerol) using a 5 mL HiTrap desalting column (GE Healthcare) and a final concentration using the ultrafiltration device described above. Final protein concentrations were determined by measuring A_280_ and using molar extinction coefficients calculated using Expasy’s ProtParam tool (https://web.expasy.org/protparam/) assuming all cysteines to be in the reduced form (ε= 1490 for AcpP; ε=28880 for Sfp).

### Sequence analysis

Sequences for Camelina ACP proteins were originally obtained from (Nguyen et al., 2013). All listed sequences are described in NCBI.

### Protein assay and SDS-PAGE

Protein Assay Reagent (Bio-Rad) was used for quantification of biological protein extractions with bovine serum albumin as the protein standard. Mini-PROTEAN tris-tricine precast gels, 4-15% and 16.5% were used for SDS-PAGE analysis. Proteins were visualized by coomassie staining, imaged with Gel Doc™ EZ and Image Lab 6.0.1 software (Bio-Rad), and reaction efficiency analyzed by densitometry using ImageJ (Fiji version 2.0.0).

### Acyl-ACP preparation

Acyl-ACPs were enriched using trichloroacetic acid (TCA) precipitation. In brief, approximately 50 mg of fresh plant tissue was powdered in the presence of liquid nitrogen. Five percent TCA was added, and the solution was vortexed. The protein was pelleted with centrifugation at 21,000 x g for 10 min at 4°C. The supernatant was removed, and the process was repeated with precipitation in 1% TCA. Next the pellet was resuspended in 50 mM 3-(N-Morpholino)propanesulfonic acid (MOPS) buffer (pH 7.5) and incubated on ice for 1 hr, then centrifuged at 21,000 x g for 10 min at 4°C to remove cellular debris. Supernatant was collected, filtered, and TCA was added to a final concentration of 10%. The sample was frozen at −80°C for 1 hr or overnight. After thawing on ice, the pellet was recovered by centrifugation at 21,000 x g for 10 min at 4°C and rinsed with 1% TCA. A final centrifugation at 21,000 x g for 10 min at 4°C ensued prior to dissolving in a minimal volume of 50 mM MOPS buffer (pH 7.5).

### Asp-N digestion

Acyl-ACP protein levels were quantified then samples were digested with endoproteinase Asp-N (Sigma) at 1:20 (enzyme:acyl-ACP, w/w), and incubated at 37°C overnight. Methanol was added to a final concentration of 50% after the enzyme digestion.

### Assessing ACP losses during initial development of the protein purification steps

To gauge individual losses of ACPs during processing steps, (though not crucial to the isotope dilution-based quantification); analyte acyl-ACPs from plant biomass were spiked with a 15:0-ACP standard before and during extraction steps as part of the initial development of the method. Recovery of a 15:0-ACP standard did not indicate a pattern in losses and was not specific to a particular step or changes in TCA concentration from 25% to 1% (Supplemental Tables S5 and S6).

### Liquid chromatography mass spectrometry condition

The Asp-N digestion products of acyl-ACPs were analyzed using a liquid chromatography tandem triple quadrupole mass spectrometer (QTRAP® 6500 LC/MS/MS system, SCIEX, Framingham, MA). Analyst software (SCIEX, version 1.6) was used to collect and analyze the data. The partial acyl-ACP molecules were first separated on a reversed-phase column (Discovery® BIO Wide Pore C18; Sigma, 10 cm, 2.1 mm, 3 μm) using solvent A (acetonitrile/10 mM ammonium formate and formic acid, pH 3.5, 10/90, v/v) and solvent B (acetonitrile/10 mM ammonium formate and formic acid, pH 3.5, 90/10, v/v). A flow rate of 0.2 ml/min was used with a gradient program: 0% B (100% A), at four minutes change to: 10% B, then to 100% B at 12 min with a hold for an additional five minutes, then changed back to 0% B at 18min and stopped at 27 min. The MS/MS spectrometry data were collected in positive ion mode. The curtain gas was set to 30 psi, the ion spray voltage was 4.5 kV, the source temperature was 400°C, the nebulizing gas (GS1) was 30 psi, and the focusing gas (GS2) was at 30 psi. The digested acyl-ACP molecules were identified using multiple-reaction monitoring (MRM) mode. The masses for the precursor ions and product ions were calculated and an MRM list was generated. Declustering potential (DP) and collision energy (CE) were optimized by changing the values stepwise with standards or plant samples. The MRM masses, DP, and CE for routine analysis are recorded in Fig. 3c and the values for fatty acid elongation intermediates including 3-hydroxyacyl-ACPs and 2,3-*trans*-enoyl-ACPs are presented in Supplemental Table S2.

### Optimization conditions for Sfp transferase reactions

Optimization of Sfp transferase reactions comprised of varying 50 mM MOPS from pH 6.5 to 7.5, DTT or TCEP from 0 to 5 mM, MgCl_2_ from 5 to 20 mM, MnCl_2_ from 5 to 20 mM, acyl+ or BODIPY-CoA from 150 to 500 μM, Sfp from 1 to 20 μM and apo-ACP from 70 to 250 μM and were performed at room, reduced (4°C), or elevated (37°C) temperature for 1 hour to 1 week.

### BODIPY-CoA chemical synthesis

Chemical synthesis of the fluorophore BODIPY-FL-*N-*(2-aminoethyl) maleimide-S-CoA (BODIPY-CoA) was accomplished by reacting 2.5 mg/mL BODIPY-FL-*N-*2-aminoethyl)maleimide with 2.5 mg/mL CoA, tri-lithium in 100 mM MES, 100 mM Mg(OAC)_2_, and 15% DMSO at 4°C in the dark for 20 minutes. The reaction efficiency was assessed by thin layer chromatography. Unreacted dye was removed by extracting with ethyl acetate three times.

### Preparation of isotopically labeled acyl-ACP standards

^15^N labeled acyl-ACP standards were synthesized enzymatically from ^15^N apo-ACP (overexpression in *E. coli*) with acyl-CoA substrates using 4’-phosphopantetheinine transferase Sfp (Sfp) from *Bacillus subtilis* overexpressed in *E. coli.* Reactions performed in 50 mM MOPS (pH 6.5), 4 mM TCEP, 100-250 μM apo-ACP, 300-500 μM acyl-CoA, 10% DMSO, 1% TWEEN-20, 10 mM MgCl_2_, 10 mM MnCl_2_, and 25 μM Sfp were incubated for 3 hours at 37°C.

Synthesized acyl-ACP standards were purified by TCA precipitation and the reaction yields were determined. TCA was added to the reaction mixture to a final concentration of 5% and the solution vortexed. The acyl-ACP was then pelleted by centrifugation at 21,000 x g at 4°C, supernatant discarded and washed with one percent TCA followed by centrifugation. Pellets were resuspended in MOPS pH 7.5 prior to use as an internal standard. The yield of each acyl-ACP standard was determined by SDS-PAGE on 16.5% tris-tricine gels and confirmed by LC-MS/MS.

### Isotope dilution-based quantitation of ACPs in biological samples

Acyl-ACPs from seed and leaf tissue were quantified using internal standards and an isotope dilution-based approach. Unlabeled standards were serially diluted with a constant concentration of ^15^N_3_ acyl-ACP standards. The area ratios of unlabeled and labeled peaks were plotted against the concentration of the unlabeled standards in order to construct standard curves. Acyl ACPs from unlabeled camelina were spiked with ^15^N_3_-labeled ACP standards then prepared and analyzed as described above with quantities calculated using the aforementioned standard curves. All peak integration was performed using Analyst 1.6 software (SCIEX). Standard curves and quantitative calculations were performed using peak areas exported to Microsoft Excel.

## ACKNOWLEDGMENTS

The authors would like to thank Chiara Bigogno for initial experiments on acyl-ACP analysis, Drs. Ed Cahoon and Jillian Silva for initial supplies of 14:0-, 15:0-, and 16:0-ACP, and Drs. John Shanklin and Ed Whittle for initial supplies of 18:1-ACP. This research was supported by the USDA-ARS, USDA-NIFA grant award (2017-67013-26156), NSF-MCB (DBI-1616820), NSF-PGRP (DBI-1828365), NSF-REU (DBI-1659812) and the Department of Energy (ARPA-e award: DE-AR0000202 and BES award: DE-SC0001295). Mass spectrometry was performed at the Donald Danforth Plant Science Center Proteomics and Mass Spectrometry Core Facility and involved QTRAP LC-MS/MS instruments acquired through National Science Foundation Major Research Instrumentation Awards (DBI-1427621) and (DBI-0521250). Any product or trademark mentioned here does not imply a warranty, guarantee, or endorsement by the authors or their affiliations over other suitable products.

## AUTHOR CONTRIBUTIONS

JGJ and DKA conceived and designed profiling experiments BSE and DKA developed quantification methods, J-WN, LMJ and JL performed experiments and analyzed data, J-WN, LMJ, BSE, JGJ and DKA wrote the manuscript that was approved by all authors.

## Suplemental Data

**Figure S1.**
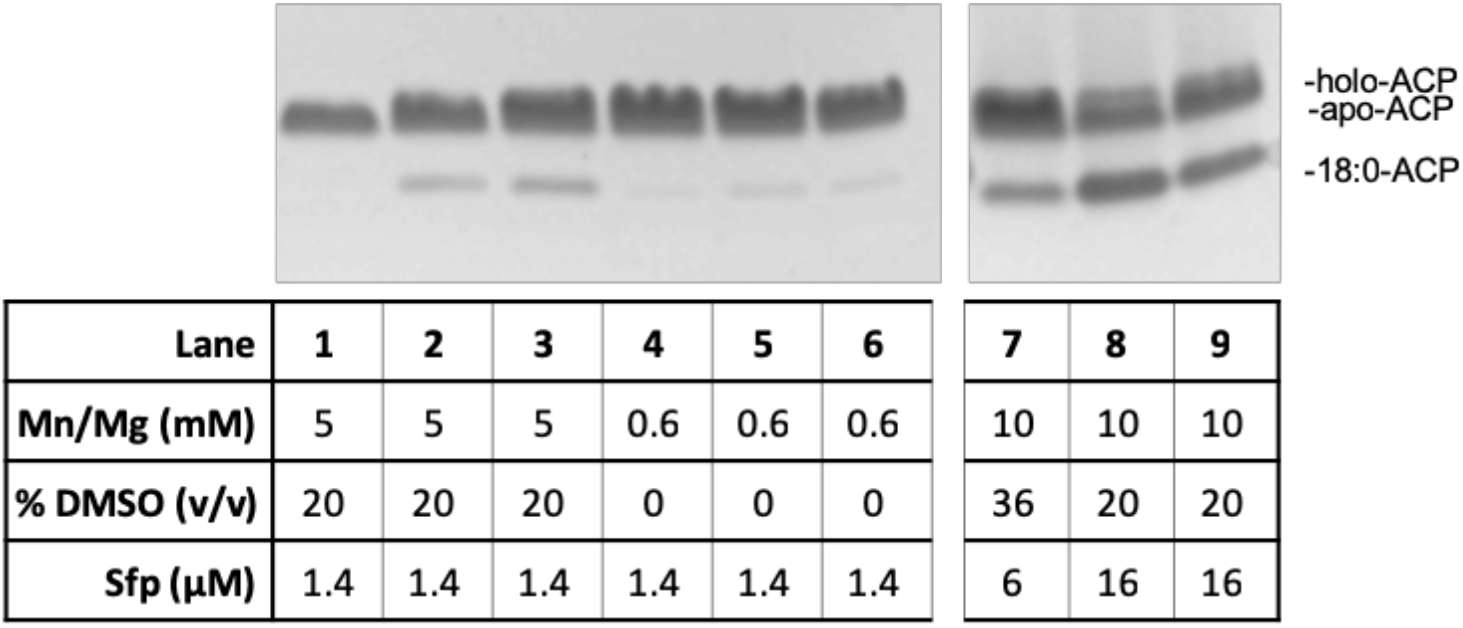
Decreased MgCl_2_ and MnCl_2_ levels decreased 18:0-ACP synthesis by Sfp. A decrease in MgCl_2_ and MnCl_2_ from 5 mM (lanes 2-3) to 0.6 mM (lanes 5-6) resulted in a decrease in 18:0-ACP production. Reactions containing 10 mM MgCl_2_ and MnCl_2_ exhibit increased production of 18:0-ACP (lanes 7-8). A portion of this increase is likely attributed to the increase in Sfp concentration, as seen in the increase in 18:0-ACP between lanes 7 and 8-9. Lanes; 1,4 unreacted apo-ACP (negative control); 2-3, 5 mM MgCl_2_ and MnCl_2_ reactions; 5-6, 0.6 mM MgCl_2_ and MnCl_2_ reactions; 7-8, 10 mM MgCl_2_ and MnCl_2_ reactions.

**Figure S2.**
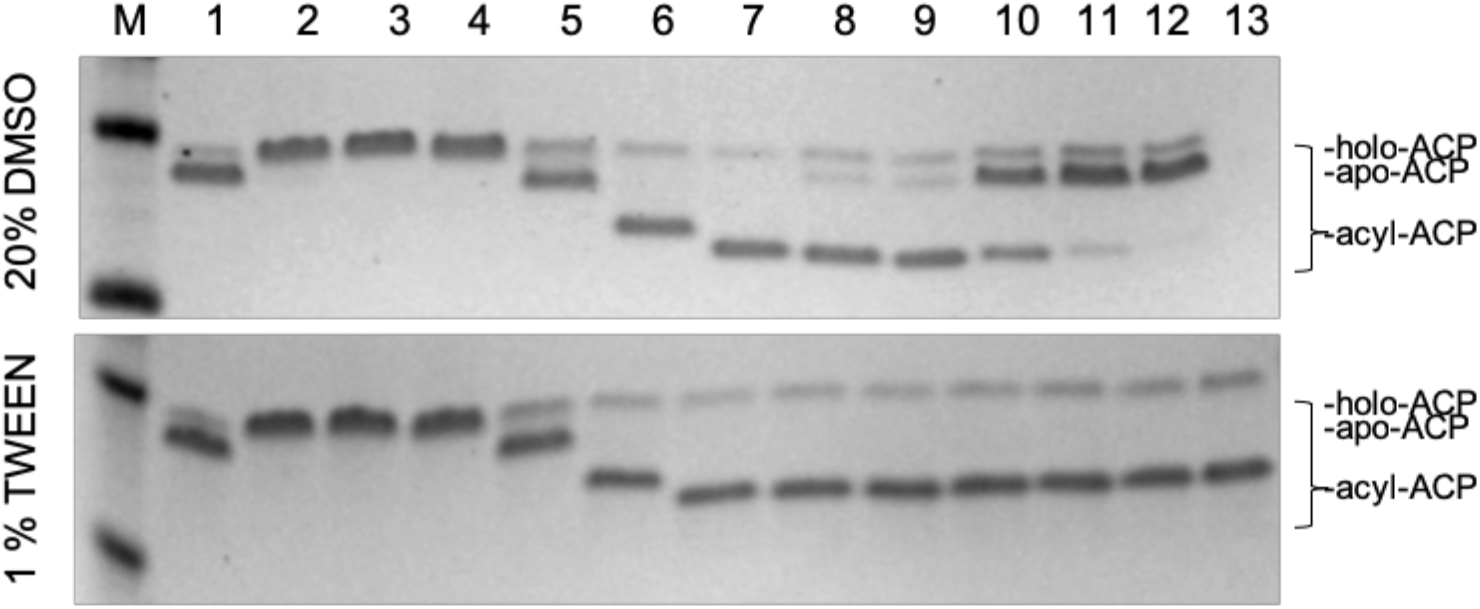
Solubilization of acyl-CoA reactants using DMSO and TWEEN-20 increases reaction efficiency. **(Top)** Inclusion of 20% DMSO (v/v) increased reaction efficiency; however, 10:0- and 12:0-ACP reactions only partially reacted and 14:0-, 16:0-, and 18:0-ACP reactions remained largely incomplete indicated by the presence of apo-ACP. **(Bottom)** Inclusion of 1% (w/v) TWEEN-20 resulted in complete reactions for all acyl-chain lengths. Holo-, malonyl-, and 2:0-ACP are indistinguishable by migration; reaction efficiency was therefore determined by the absence of the apo-ACP substrate band. Lanes; M, marker; 1, apo-ACP (no reaction); 2, holo-ACP; 3, malonyl-ACP; 4, 2:0-ACP; 5, 4:0-ACP; 6, 6:0-ACP; 7, 8:0-ACP; 8, 10:0-ACP; 9, 12:0-ACP; 10, 14:0-ACP; 11, 16:0-ACP; 12, 18:0-ACP; 13, 18:1-ACP (bottom only).

**Figure S3.**
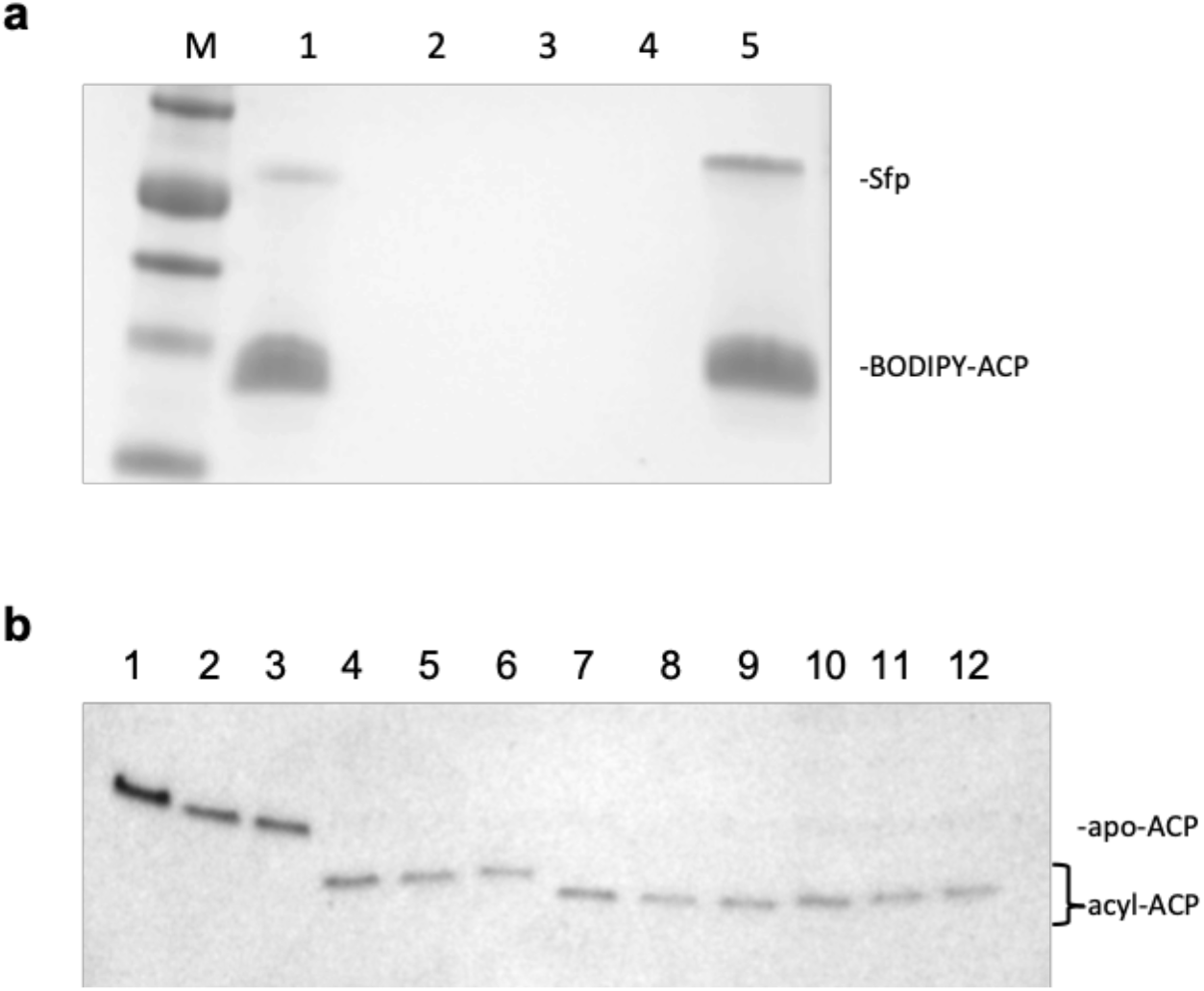
Acyl-ACP Standards are quantitatively recovered by trichloroacetic acid (TCA). **a)** No losses were observed in the 5% or 1% TCA wash steps (lanes 2-4), and quantitative recovery of BODIPY-ACP was achieved (lanes 1 and 5). Lanes; 1, BODIPY-ACP (before extraction); 2, 5% (w/v) TCA supernatant; 3, 1% (w/v) TCA wash; 4, 1% (w/v) TCA supernatant; 5, 50 mM MOPS, pH 7.5 resuspension. **b)** TCA recovery of acyl-ACP standards. Lanes; 1, apo-ACP (before extraction); 2-3 apo-ACP (after extraction); 4, 6:0-ACP (before extraction); 5-6 6:0-ACP (after extraction); 7, 14:0-ACP (before extraction); 8-9 14:0-ACP (after extraction); 10, 18:0-ACP (before extraction); 11-12 18:0-ACP (after extraction).

**Figure S4.**
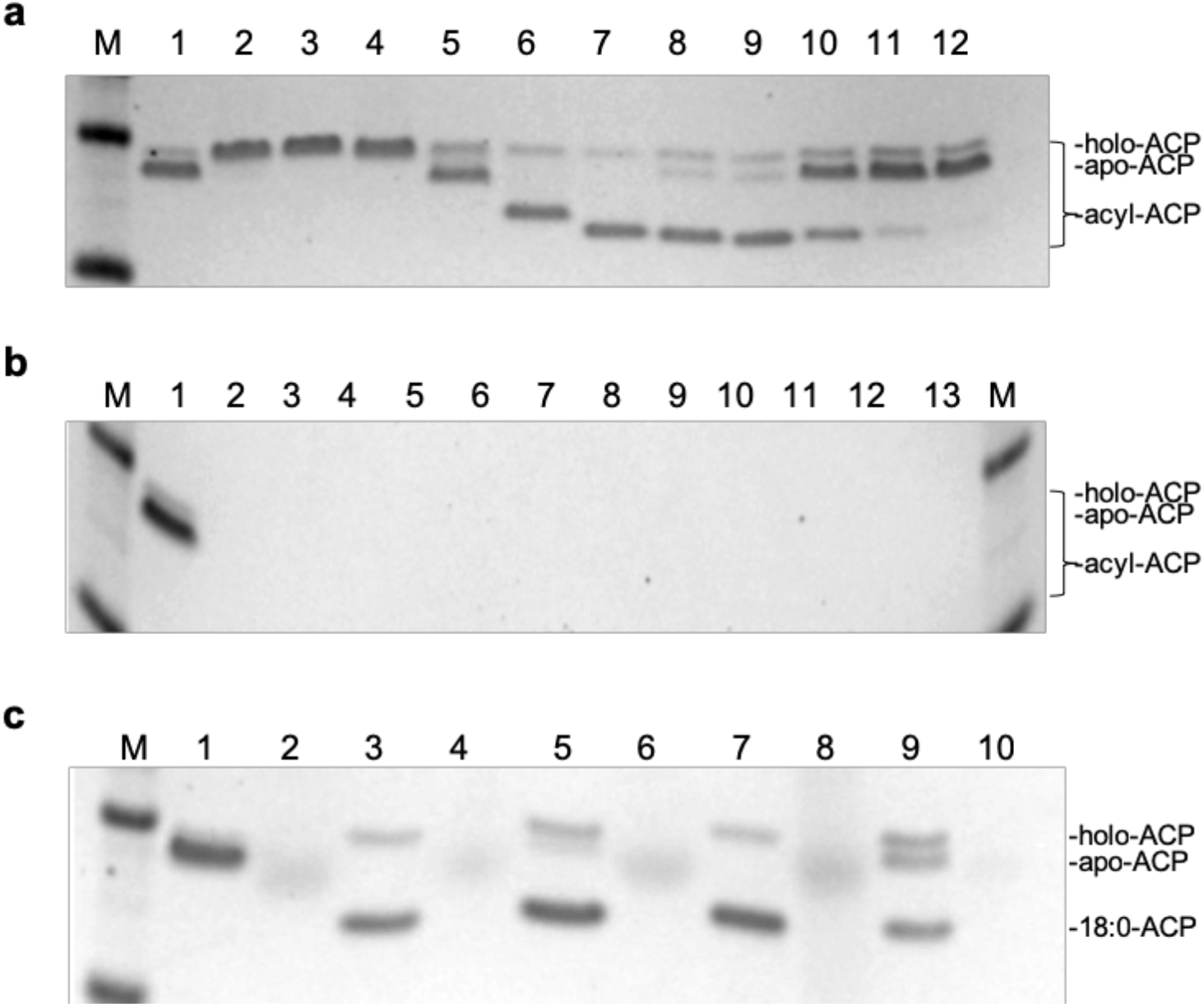
Complete digestion of acyl-ACP standards. **a)** SDS-PAGE analysis of acyl-ACP standards before asp-N (*image is duplicated from Fig S2 for ease of comparison here*). Lanes; M, marker; 1, apo-ACP; 2, holo-ACP; 3, malonyl-ACP; 4, 2:0-ACP; 5, 4:0-ACP; 6, 6:0-ACP; 7, 8:0-ACP; 8, 10:0-ACP; 9, 12:0-ACP; 10, 14:0-ACP; 11, 16:0-ACP; 12, 18:0-ACP. **b)** SDS-PAGE analysis of acyl-ACP standards after asp-N. All ACPs are digested by 1:50 and 1:20 enzyme:protein for apo-through 8:0-ACP and 10:0- through 18:0-ACP, respectively. Lanes; M, marker; 1, apo-ACP (undigested); 2, apo-ACP (digested); 3, holo-ACP (digested); 4, malonyl-ACP (digested); 5, 2:0-ACP (digested); 6, 4:0-ACP (digested); 7, 6:0-ACP (digested); 8, 8:0-ACP (digested); 9, 10:0-ACP (digested); 10, 12:0-ACP (digested); 11, 14:0-ACP (digested); 12, 16:0-ACP (digested); 13, 18:0-ACP (digested). **c)** SDS-PAGE analysis of 18:0-ACP standards before and after asp-N. Lanes; 1, apo-ACP (undigested); 2, apo-ACP (digested); 3, 5, 7, 9, 18:0-ACP (undigested); 4, 6, 8, 10, 18:0-ACP (digested).

**Figure S5.**
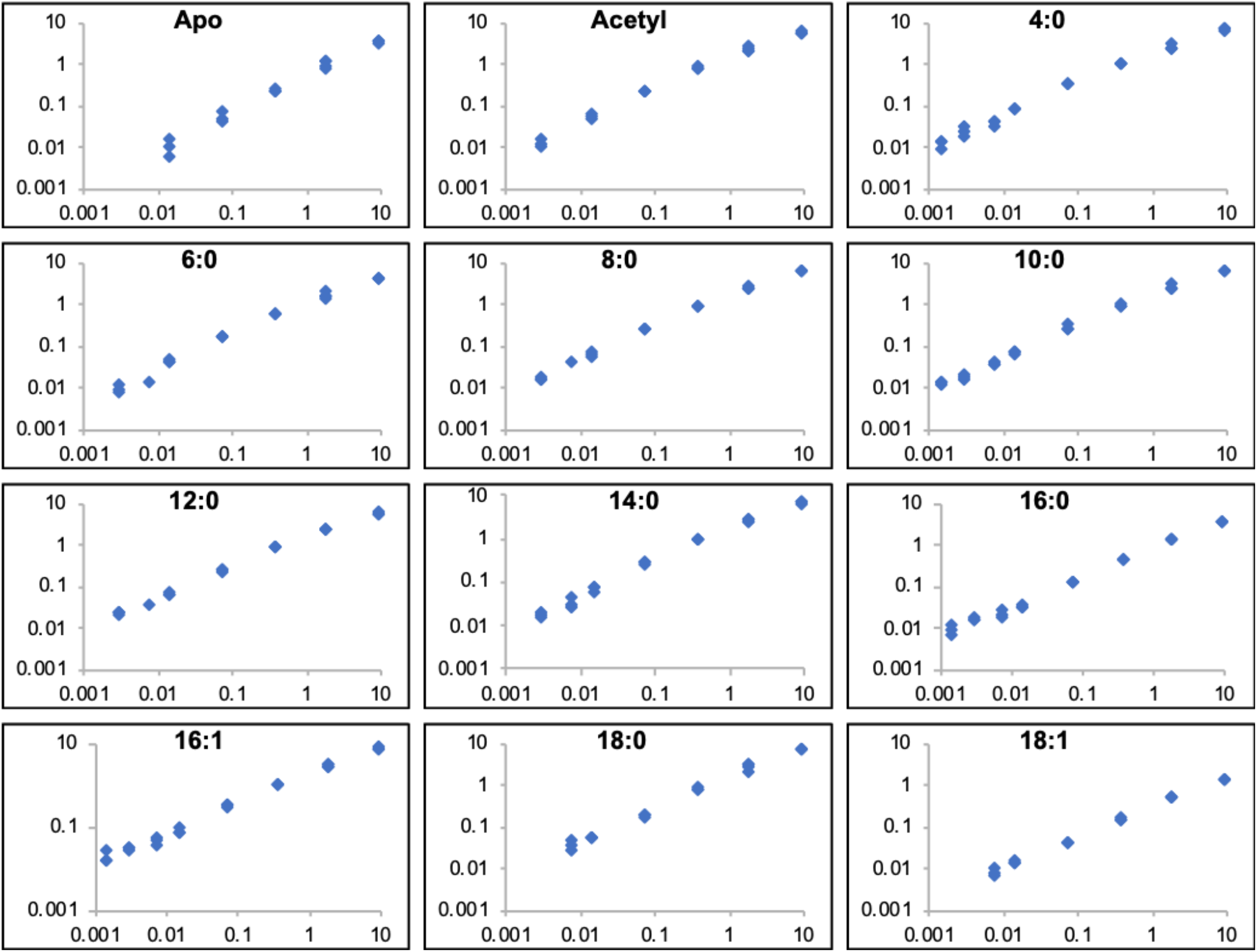
Calibration curves of acyl-ACPs. Acyl-ACPs were measured over a range of concentrations to establish the linearity of response, limit of quantification and dynamic range (n=3). Data are presented on logarithmic scales. Y-axes, peak area analyte/peak area standard; x-axes, pmol analyte. See Figure 3 in the main text for further details.

**Figure S6.**
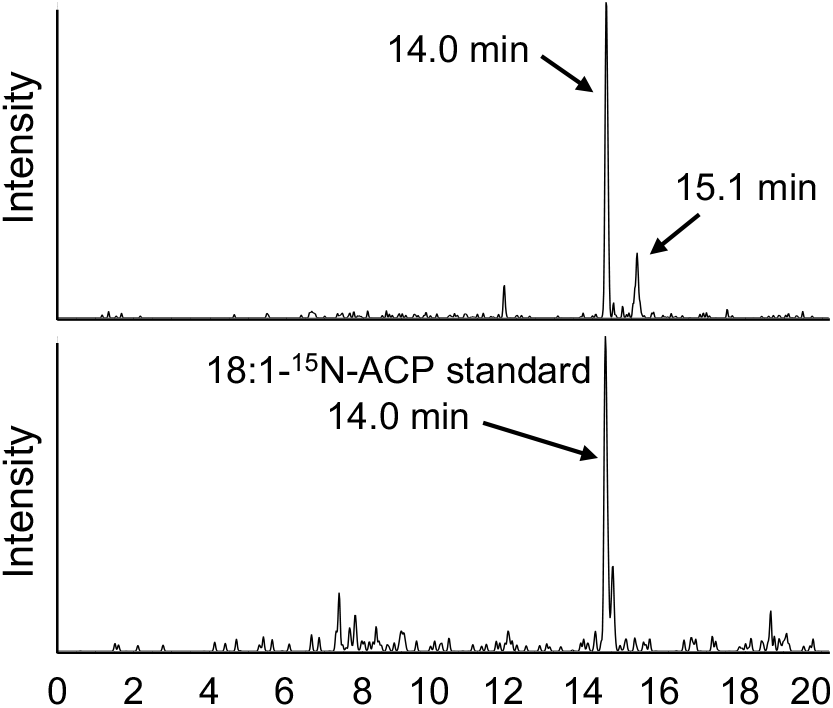
Retention time for the 18:1-ACP elongation intermediate (i.e. C18-enoyl-ACP or 2,3-*trans*-octadecenoyl-ACP). 18:1-ACP (i.e. 9-cis-18:1-ACP) and the C18-enoyl-ACP isomer of 18:1 have the same molecular composition and therefore cannot be differentiated by mass alone as indicated in the upper panel that is a chromatogram extraction of the shared *m/z*. An isotopically labeled internal standard containing predominantly 18:1-ACP was used to assign the retention time for oleoyl- and the C18-enoyl-ACP isomer. As expected, the much smaller peak in the upper panel was the C18-enoyl-ACP isomer that is lower in concentration. Axes were redrawn for clarity.

**Figure S7.**
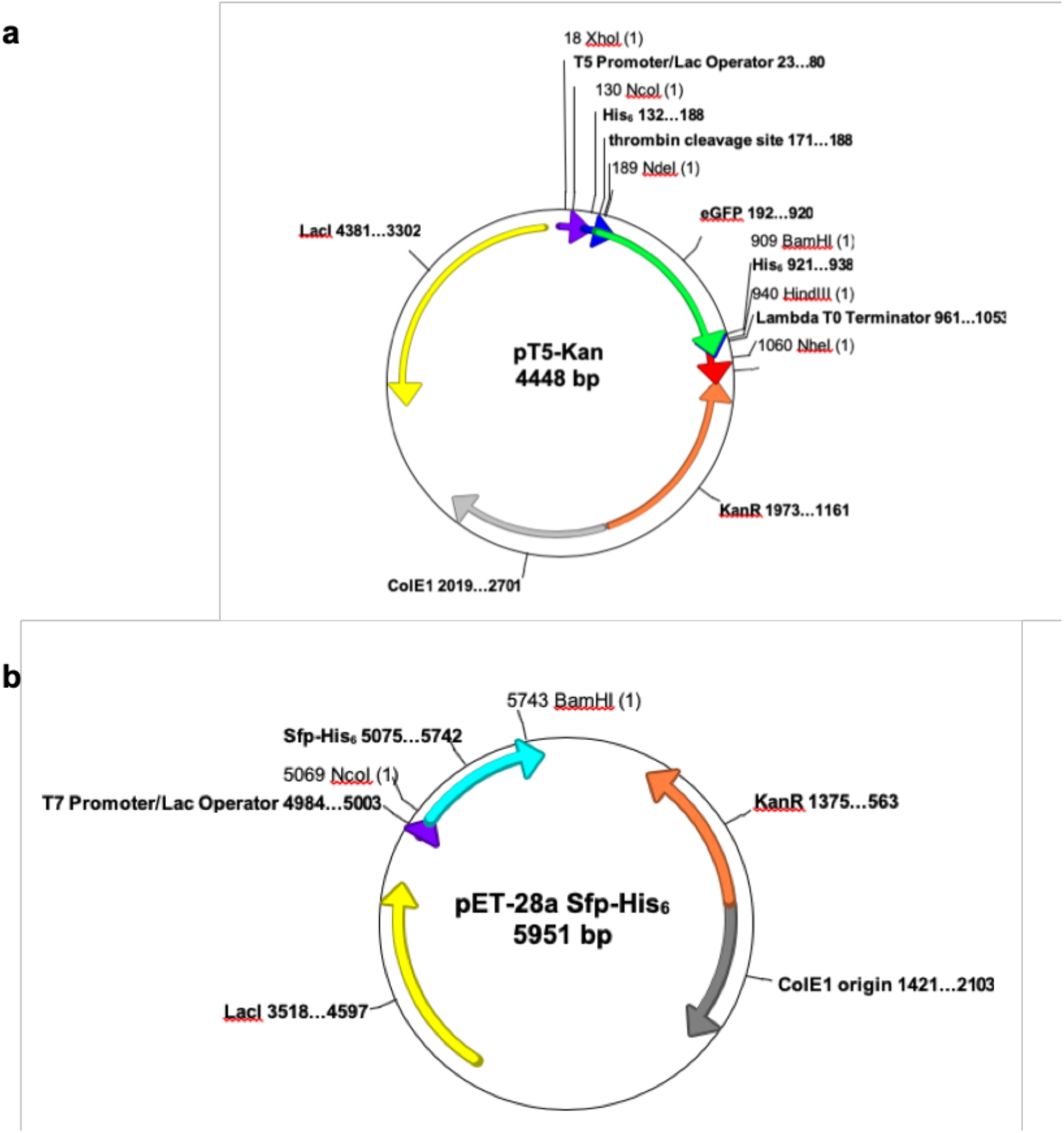
Plasmid design for ACP and Sfp transferase overexpression. **a)** Plasmid pT5Kan. **b)** Plasmid pET-28a-Sfp-His_6_. The ColEI origin maintains high copy number plasmids. Plasmids contained restriction sites that allowed for expression of proteins with either N-terminal (removable via thrombin cleavage) and/or C-terminal hexahistidine tags. Expression was repressed under noninducing conditions by tandem *lac* operator sites recognized by LacI repressor (encoded by the plasmid) binding sites present in the T5/T7 promoter. Induction of expression was achieved using IPTG, lactose or standard autoinduction protocols. Selection is made possible by a kanamycin resistance gene encoded by the plasmid (aminoglycoside-O-phosphotransferase).

**Table S1.**
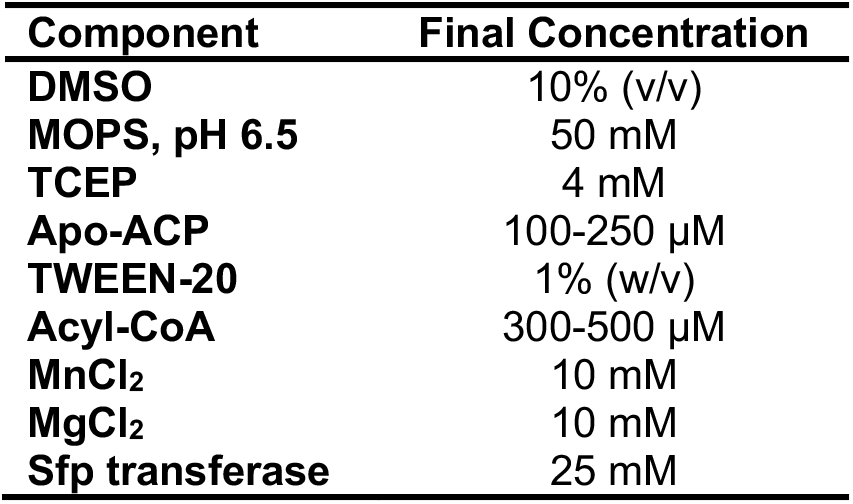
Acyl-ACP synthesis reaction. Components were added in the order listed with adequate mixing before addition of MnCl_2_ and MgCl_2_ to avoid substrate precipitation.

**Table S2.** MRM list for acyl-ACP elongation intermediates. Masses based on composition and expected fragmentation patterns were used to examine the presence of acyl-ACP elongation intermediates. The optimized declustering potential (DP) and collision energy (CE) used were the same as described in Figure 3C.

| Intermediate Acyl (-ACP) | Chemical Formula | M+H | -413.1 | DP, CE, CXP |
| --- | --- | --- | --- | --- |
| Acetyl | C<sub>2</sub>H<sub>3</sub>O | 716.3 | 303.2 | 80, 40, 10 |
| 3-Ketobutyryl | C<sub>4</sub>H<sub>5</sub>O<sub>2</sub> | 758.3 | 345.2 | 90, 50, 10 |
| 3-Hydroxybutyryl | C<sub>4</sub>H<sub>7</sub>O<sub>2</sub> | 760.3 | 347.2 | 90, 50, 10 |
| 2,3-trans-Butenoyl | C<sub>4</sub>H<sub>5</sub>O | 742.3 | 329.2 | 90, 50, 10 |
| Butyryl (4:0) | C<sub>4</sub>H<sub>7</sub>O | 744.3 | 331.2 | 90, 40, 10 |
| 3-Ketohexanoyl | C<sub>6</sub>H<sub>9</sub>O<sub>2</sub> | 786.3 | 373.2 | 100, 50, 10 |
| 3-Hydroxyhexanoyl | C<sub>6</sub>H<sub>11</sub>O<sub>2</sub> | 788.3 | 375.2 | 100, 50, 10 |
| 2,3-trans-Hexenoyl | C<sub>6</sub>H<sub>9</sub>O | 770.3 | 357.2 | 100, 50, 10 |
| Hexanoyl (6:0) | C<sub>6</sub>H<sub>11</sub>O | 772.3 | 359.2 | 100, 40, 10 |
| 3-Ketooctanoyl | C<sub>8</sub>H<sub>13</sub>O<sub>2</sub> | 814.3 | 401.2 | 100, 50, 10 |
| 3-Hydroxyoctanoyl | C<sub>8</sub>H<sub>15</sub>O<sub>2</sub> | 816.3 | 403.2 | 100, 45, 10 |
| 2,3-trans-Octenoyl | C<sub>8</sub>H<sub>13</sub>O | 798.3 | 385.2 | 100, 40, 10 |
| Octanoyl (8:0) | C<sub>8</sub>H<sub>15</sub>O | 800.4 | 387.3 | 100, 45, 10 |
| 3-Ketodecanoyl | C<sub>10</sub>H<sub>17</sub>O<sub>2</sub> | 842.4 | 429.3 | 100, 40, 10 |
| 3-Hydroxydecanoyl | C<sub>10</sub>H<sub>19</sub>O<sub>2</sub> | 844.4 | 431.3 | 100, 40, 10 |
| 2,3-trans-Decenoyl | C<sub>10</sub>H<sub>17</sub>O | 826.4 | 413.3 | 100, 40, 10 |
| Decanoyl (10:0) | C<sub>10</sub>H<sub>19</sub>O | 828.4 | 415.3 | 100, 45, 10 |
| 3-Ketododecanoyl | C<sub>12</sub>H<sub>21</sub>O<sub>2</sub> | 870.4 | 457.3 | 100, 40, 10 |
| 3-Hydroxydodecanoyl | C<sub>12</sub>H<sub>23</sub>O<sub>2</sub> | 872.4 | 459.3 | 100, 40, 10 |
| 2,3-trans-Dodecenoyl | C<sub>12</sub>H<sub>21</sub>O | 854.4 | 441.3 | 100, 40, 10 |
| Dodecanoyl (12:0) | C<sub>12</sub>H<sub>23</sub>O | 856.4 | 443.3 | 100, 45, 10 |
| 3-Ketotetradecanoyl | C<sub>14</sub>H<sub>25</sub>O<sub>2</sub> | 898.4 | 485.3 | 100, 45, 10 |
| 3-Hydroxytetradecanoyl | C<sub>14</sub>H<sub>27</sub>O<sub>2</sub> | 900.4 | 487.3 | 100, 45, 10 |
| 2,3-trans-Tetradecenoyl | C<sub>14</sub>H<sub>25</sub>O | 882.4 | 469.3 | 100, 45, 10 |
| Tetradecanoyl (14:0) | C<sub>14</sub>H<sub>27</sub>O | 884.4 | 471.3 | 100, 45, 10 |
| 3-Ketohexadecanoyl | C<sub>16</sub>H<sub>29</sub>O<sub>2</sub> | 926.5 | 513.4 | 100, 45, 10 |
| 3-Hydroxyhexadecanoyl | C<sub>16</sub>H<sub>31</sub>O<sub>2</sub> | 928.5 | 515.4 | 100, 45, 10 |
| 2,3-trans-Hexadecenoyl | C<sub>16</sub>H<sub>29</sub>O | 910.5 | 497.4 | 100, 45, 10 |
| Hexadecanoyl (16:0) | C<sub>16</sub>H<sub>31</sub>O | 912.5 | 499.4 | 100, 50, 10 |
| cis, cis, cis-9,12,15-hexadecanoyl (16:3) | C<sub>16</sub>H<sub>25</sub>O | 906.4 | 493.3 | 85,46,14 |
| 3-Ketooctadecanoyl | C<sub>18</sub>H<sub>33</sub>O<sub>2</sub> | 954.5 | 541.4 | 100, 45, 10 |
| 3-Hydroxyoctadecanoyl | C<sub>18</sub>H<sub>35</sub>O<sub>2</sub> | 956.5 | 543.4 | 100, 45, 10 |
| 2,3-trans-Octadecenoyl | C<sub>18</sub>H<sub>33</sub>O | 938.5 | 525.4 | 100, 45, 10 |
| Octadecanoyl (18:0) | C<sub>18</sub>H<sub>35</sub>O | 940.5 | 527.4 | 100, 50, 10 |

**Table S3.** Retention time of acyl-ACP elongation intermediates of the fatty acid biosynthetic cycle. Relative retention times for the intermediates of the fatty acid biosynthetic cycle were inspected. The ketoacyl-ACP intermediate was not identified.

| Chain length | Relative retention time (min) |  |
| --- | --- | --- |
|  | 3-hydroxyacyl-ACP | 2,3-trans-enoyl-ACP |
| C4 | 6.7 | 8.0 |
| C6 | 7.9 | 8.9 |
| C8 | 8.9 | 9.8 |
| C10 | 9.8 | 10.7 |
| C12 | 10.7 | 11.7 |
| C14 | 11.7 | 12.8 |
| C16 | 12.8 | 14.0 |
| C18 | 14.0 | 15.1 |

**Table S4.**
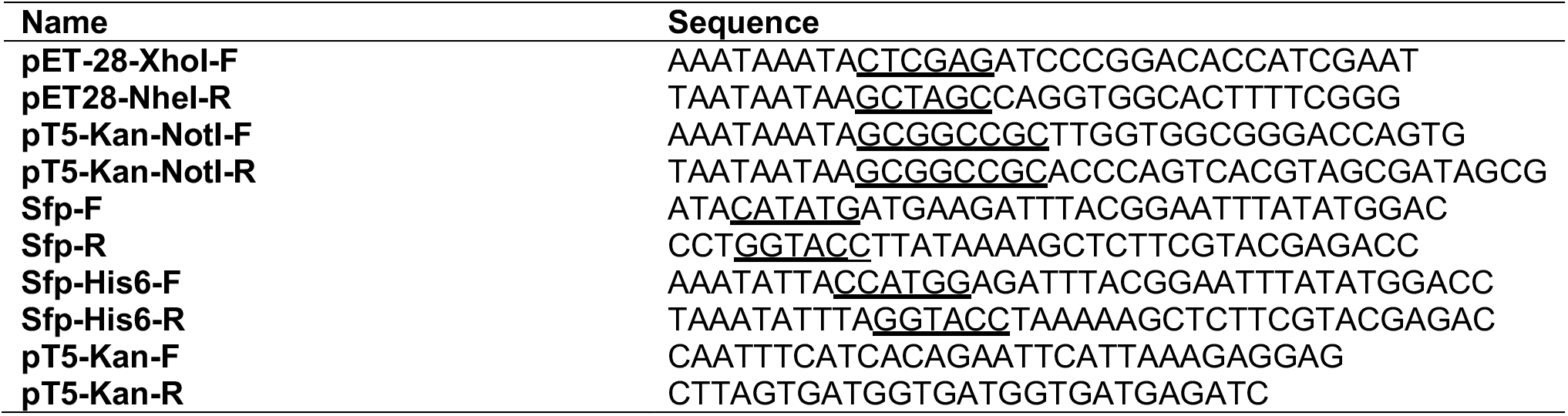
Primers used for plasmid construction and sequencing. Restriction sites are underlined.

**Table S5.** 15:0-ACP recovery. A 15:0-ACP internal standard was added at different points in the protein extraction to ensure reasonable recovery during the development of the extraction procedure. Each peak area is the average of two replicates and indicates that there are not major losses at a particular step in the protein extraction protocol. Importantly, the isotope dilution approach accounts for any losses in a sample because of the presence of internal standards; however, the initial 15:0-ACP recovery here was used to help develop the method prior to absolute quantification.

| **Description of 15:0 addition** | **Peak area** |
| --- | --- |
| **With biomass** | 488,800 |
|  | 512,000 |
| **With 1% TCA** | 424,950 |
|  | 410,500 |
| **With MOPS** | 571,700 |
|  | 398,900 |
| **With 2<sup>nd</sup> MOPS** | 349,450 |
|  | 372,450 |
| **Prior to ASP-N** | 687,400 |
|  | 463,450 |

**Table S6.** Recovery of 15:0-ACP with different TCA concentrations. A 15:0-ACP internal standard was precipitated with different amounts of TCA, resulting in similar peak areas regardless of the TCA addition.

| **Amount of**<br>**TCA (w/v)** | **Peak area** |
| --- | --- |
| **25%** | 3,209,000 |
| **20%** | 3,891,000 |
| **15%** | 4,008,000 |
| **10%** | 3,778,000 |
| **5%** | 3,952,000 |
| **1%** | 3,387,000 |

## Parsed Citations

Alberts, A.W., Goldman, P., and Vagelos, P.R. (1963). The condensation reaction of fatty acid synthesis. I. Separation and properties of the enzymes. The Journal of biological chemistry 238, 557–565. Pubmed: Author and Title Google Scholar: Author Only Title Only Author and Title

Allen, D.K. (2016a). Assessing compartmentalized flux in lipid metabolism with isotopes. Biochimica et biophysica acta 1861, 1226–1242. Pubmed: Author and Title Google Scholar: Author Only Title Only Author and Title

Allen, D.K. (2016b). Quantifying plant phenotypes with isotopic labeling & metabolic flux analysis. Curr Opin Biotechnol 37, 45–52. Pubmed: Author and Title Google Scholar: Author Only Title Only Author and Title

Andre, C., Haslam, R.P., and Shanklin, J. (2012). Feedback regulation of plastidic acetyl-CoA carboxylase by 18:1-acyl carrier protein in Brassica napus. Proceedings of the National Academy of Sciences of the United States of America 109, 10107–10112. Pubmed: Author and Title Google Scholar: Author Only Title Only Author and Title

Bonaventure, G., and Ohlrogge, J.B. (2002). Differential regulation of mRNA levels of acyl carrier protein isoforms in Arabidopsis. Plant physiology 128, 223–235. Pubmed: Author and Title Google Scholar: Author Only Title Only Author and Title

Bressler, R., and Wakil, S.J. (1961). Studies on the Mechanism of Fatty Acid Synthesis: IX. The conversion of malonyl coenzyme A to long chain fatty acids. The Journal of biological chemistry 236, 1643–1651. Pubmed: Author and Title Google Scholar: Author Only Title Only Author and Title

Brody, S., Oh, C., Hoja, U., and Schweizer, E. (1997). Mitochondrial acyl carrier protein is involved in lipoic acid synthesis in Saccharomyces cerevisiae. FEBS letters 408, 217–220. Pubmed: Author and Title Google Scholar: Author Only Title Only Author and Title

Chuman, L., and Brody, S. (1989). Acyl carrier protein is present in the mitochondria of plants and eucaryotic micro-organisms. European journal of biochemistry / FEBS 184, 643–649. Pubmed: Author and Title Google Scholar: Author Only Title Only Author and Title

Constantinides, P.P., and Steim, J.M. (1986). Solubility of palmitoyl-coenzyme Ain acyltransferase assay buffers containing magnesium ions. Archives of biochemistry and biophysics 250, 267–270. Pubmed: Author and Title Google Scholar: Author Only Title Only Author and Title

Cronan, J.E. (2014). The chain-flipping mechanism of ACP (acyl carrier protein)-dependent enzymes appears universal. Biochem J 460, 157–163. Pubmed: Author and Title Google Scholar: Author Only Title Only Author and Title

Cronan, J.E., Fearnley, I.M., and Walker, J.E. (2005). Mammalian mitochondria contain a soluble acyl carrier protein. FEBS letters 579, 4892–4896. Pubmed: Author and Title Google Scholar: Author Only Title Only Author and Title

De Lay, N.R., and Cronan, J.E. (2007). In vivo functional analyses of the type II acyl carrier proteins of fatty acid biosynthesis. The Journal of biological chemistry 282, 20319–20328. Pubmed: Author and Title Google Scholar: Author Only Title Only Author and Title

Fan, J., Yan, C., Roston, R., Shanklin, J., and Xu, C. (2014). Arabidopsis lipins, PDAT1 acyltransferase, and SDP1 triacylglycerol lipase synergistically direct fatty acids toward beta-oxidation, thereby maintaining membrane lipid homeostasis. The Plant cell 26, 4119–4134. Pubmed: Author and Title Google Scholar: Author Only Title Only Author and Title

Goldman, P., Alberts, A.W., and Vagelos, P.R. (1963). The Condensation Reaction of Fatty Acid Synthesis: III. Identification of the protein-bound product of the reaction and its conversion to long chain fatty acids. The Journal of biological chemistry 238, 3579–3583. Pubmed: Author and Title Google Scholar: Author Only Title Only Author and Title

Hloušek-Radojčić, A., Post-Beittenmiller, D., and Ohlrogge, J.B. (1992). Expression of Constitutive and Tissue-Specific Acyl Carrier Protein Isoforms in Arabidopsis. Plant physiology 98, 206–214. Pubmed: Author and Title Google Scholar: Author Only Title Only Author and Title

Hsu, R.Y., Wasson, G., and Porter, J.W. (1965). The Purification and Properties of the Fatty Acid Synthetase of Pigeon Liver. The Journal of biological chemistry 240, 3736–3746. Pubmed: Author and Title Google Scholar: Author Only Title Only Author and Title

Jaworski, J.G., Post-Beittenmiller, D., and Ohlrogge, J.B. (1993). Acetyl-acyl carrier protein is not a major intermediate in fatty acid biosynthesis in spinach. European journal of biochemistry / FEBS 213, 981–987. Pubmed: Author and Title Google Scholar: Author Only Title Only Author and Title

Joshi, A.K., Zhang, L., Rangan, V.S., and Smith, S. (2003). Cloning, expression, and characterization of a human 4’-phosphopantetheinyl transferase with broad substrate specificity. The Journal of biological chemistry 278, 33142–33149. Pubmed: Author and Title Google Scholar: Author Only Title Only Author and Title

Kopka, J., Ohlrogge, J.B., and Jaworski, J.G. (1995). Analysis of in vivo levels of acyl-thioesters with gas chromatography/mass spectrometry of the butylamide derivative. Analytical biochemistry 224, 51–60. Pubmed: Author and Title Google Scholar: Author Only Title Only Author and Title

Lambalot, R.H., Gehring, A.M., Flugel, R.S., Zuber, P., LaCelle, M., Marahiel, M.A., Reid, R., Khosla, C., and Walsh, C.T. (1996). Anew enzyme superfamily - the phosphopantetheinyl transferases. Chem Biol 3, 923–936. Pubmed: Author and Title Google Scholar: Author Only Title Only Author and Title

Li-Beisson, Y., Shorrosh, B., Beisson, F., Andersson, M.X., Arondel, V., Bates, P.D., Baud, S., Bird, D., Debono, A., Durrett, T.P., Franke, R.B., Graham, I.A., Katayama, K., Kelly, A.A., Larson, T., Markham, J.E., Miquel, M., Molina, I., Nishida, I., Rowland, O., Samuels, L., Schmid, K.M., Wada, H., Welti, R., Xu, C., Zallot, R., and Ohlrogge, J. (2013). Acyl-lipid metabolism. In Arabidopsis Book, pp. e0161. Pubmed: Author and Title Google Scholar: Author Only Title Only Author and Title

Li, C., Guan, Z., Liu, D., and Raetz, C.R. (2011). Pathway for lipid Abiosynthesis in Arabidopsis thaliana resembling that of Escherichia coli. Proceedings of the National Academy of Sciences of the United States of America 108, 11387–11392. Pubmed: Author and Title Google Scholar: Author Only Title Only Author and Title

Nguyen, H.T., Silva, J.E., Podicheti, R., Macrander, J., Yang, W., Nazarenus, T.J., Nam, J.W., Jaworski, J.G., Lu, C., Scheffler, B.E., Mockaitis, K., and Cahoon, E.B. (2013). Camelina seed transcriptome: a tool for meal and oil improvement and translational research. Plant biotechnology journal 11, 759–769. Pubmed: Author and Title Google Scholar: Author Only Title Only Author and Title

Ohlrogge, J.B., and Kuo, T.M. (1985). Plants have isoforms for acyl carrier protein that are expressed differently in different tissues. The Journal of biological chemistry 260, 8032–8037. Pubmed: Author and Title Google Scholar: Author Only Title Only Author and Title

Ohlrogge, J.B., Kuhn, D.N., and Stumpf, P.K. (1979). Subcellular localization of acyl carrier protein in leaf protoplasts of Spinacia oleracea. Proceedings of the National Academy of Sciences of the United States of America 76, 1194–1198. Pubmed: Author and Title Google Scholar: Author Only Title Only Author and Title

Overath, P., and Stumpf, P.K. (1964). Fat Metabolism in Higher Plants. 23. Properties of a Soluble Fatty Acid Synthetase from Avocado Mesocarp. The Journal of biological chemistry 239, 4103–4110. Pubmed: Author and Title Google Scholar: Author Only Title Only Author and Title

Peralta-Yahya, P.P., Zhang, F., del Cardayre, S.B., and Keasling, J.D. (2012). Microbial engineering for the production of advanced biofuels. Nature 488, 320–328. Pubmed: Author and Title Google Scholar: Author Only Title Only Author and Title

Post-Beittenmiller, D., Jaworski, J.G., and Ohlrogge, J.B. (1991). In vivo pools of free and acylated acyl carrier proteins in spinach. Evidence for sites of regulation of fatty acid biosynthesis. The Journal of biological chemistry 266, 1858–1865. Pubmed: Author and Title Google Scholar: Author Only Title Only Author and Title

Quadri, L.E., Weinreb, P.H., Lei, M., Nakano, M.M., Zuber, P., and Walsh, C.T. (1998). Characterization of Sfp, a Bacillus subtilis phosphopantetheinyl transferase for peptidyl carrier protein domains in peptide synthetases. Biochemistry 37, 1585–1595. Pubmed: Author and Title Google Scholar: Author Only Title Only Author and Title

Sanchez, C., Du, L., Edwards, D.J., Toney, M.D., and Shen, B. (2001). Cloning and characterization of a phosphopantetheinyl transferase from Streptomyces verticillus ATCC15003, the producer of the hybrid peptide-polyketide antitumor drug bleomycin. Chem Biol 8, 725–738. Pubmed: Author and Title Google Scholar: Author Only Title Only Author and Title

Uauy, R., Mena, P., and Rojas, C. (2000). Essential fatty acids in early life: structural and functional role. Proc Nutr Soc 59, 3–15. Pubmed: Author and Title Google Scholar: Author Only Title Only Author and Title

Vanhercke, T., Dyer, J.M., Mullen, R.T., Kilaru, A., Rahman, M.M., Petrie, J.R., Green, A.G., Yurchenko, O., and Singh, S.P. (2019). Metabolic engineering for enhanced oil in biomass. Progress in lipid research 74, 103–129. Pubmed: Author and Title Google Scholar: Author Only Title Only Author and Title

Wada, H., Shintani, D., and Ohlrogge, J. (1997). Why do mitochondria synthesize fatty acids? Evidence for involvement in lipoic acid production. Proceedings of the National Academy of Sciences of the United States of America 94, 1591–1596. Pubmed: Author and Title Google Scholar: Author Only Title Only Author and Title

White, S.W., Zheng, J., Zhang, Y.M., and Rock. (2005). The structural biology of type II fatty acid biosynthesis. Annu Rev Biochem 74, 791–831. Pubmed: Author and Title Google Scholar: Author Only Title Only Author and Title

Witkowski, A., Joshi, A.K., and Smith, S. (2007). Coupling of the de novo fatty acid biosynthesis and lipoylation pathways in mammalian mitochondria. The Journal of biological chemistry 282, 14178–14185. Pubmed: Author and Title Google Scholar: Author Only Title Only Author and Title

Zhu, L., Zou, Q., Cao, X., and Cronan, J.E. (2019). Enterococcus faecalis Encodes an Atypical Auxiliary Acyl Carrier Protein Required for Efficient Regulation of Fatty Acid Synthesis by Exogenous Fatty Acids. MBio 10. Pubmed: Author and Title Google Scholar: Author Only Title Only Author and Title

Zhu, L.H., Krens, F., Smith, M.A., Li, X., Qi, W., van Loo, E.N., Iven, T., Feussner, I., Nazarenus, T.J., Huai, D., Taylor, D.C., Zhou, X.R., Green, A.G., Shockey, J., Klasson, K.T., Mullen, R.T., Huang, B., Dyer, J.M., and Cahoon, E.B. (2016). Dedicated Industrial Oilseed Crops as Metabolic Engineering Platforms for Sustainable Industrial Feedstock Production. Sci Rep 6, 22181. Pubmed: Author and Title Google Scholar: Author Only Title Only Author and Title

Zornetzer, G.A., White, R.D., Markley, J.L., and Fox, B.G. (2006). Preparation of isotopically labeled spinach acyl-acyl carrier protein for NMR structural studies. Protein Expr Purif 46, 446–455. Pubmed: Author and Title Google Scholar: Author Only Title Only Author and Title

